# Differences in functional connectivity along the anterior-posterior axis of human hippocampal subfields

**DOI:** 10.1101/410720

**Authors:** Marshall A. Dalton, Cornelia McCormick, Eleanor A. Maguire

**Affiliations:** Wellcome Centre for Human Neuroimaging, Queen Square Institute of Neurology, University College London, UK

**Keywords:** hippocampus, hippocampal subfields, functional connectivity, pre/parasubiculum, uncus

## Abstract

There is a paucity of information about how human hippocampal subfields are functionally connected to each other and to neighbouring extra-hippocampal cortices. In particular, little is known about whether patterns of functional connectivity (FC) differ down the anterior-posterior axis of each subfield. Here, using high resolution structural MRI we delineated the hippocampal subfields in healthy young adults. This included the CA fields, separating DG/CA4 from CA3, separating the pre/parasubiculum from the subiculum, and also segmenting the uncus. We then used high resolution resting state functional MRI to interrogate FC. We first analysed the FC of each hippocampal subfield in its entirety, in terms of FC with other subfields and with the neighbouring regions, namely entorhinal, perirhinal, posterior parahippocampal and retrosplenial cortices. Next, we analysed FC for different portions of each hippocampal subfield along its anterior-posterior axis, in terms of FC between different parts of a subfield, FC with other subfield portions, and FC of each subfield portion with the neighbouring cortical regions of interest. We found that intrinsic functional connectivity between the subfields aligned generally with the tri-synaptic circuit but also extended beyond it. Our findings also revealed that patterns of functional connectivity between the subfields and neighbouring cortical areas differed markedly along the anterior-posterior axis of each hippocampal subfield. Overall, these results contribute to ongoing efforts to characterise human hippocampal subfield connectivity, with implications for understanding hippocampal function.

**Highlights:**

- High resolution resting state functional MRI scans were collected
- We investigated functional connectivity (FC) of human hippocampal subfields
- We specifically examined FC along the anterior-posterior axis of subfields
- FC between subfields extended beyond the canonical tri-synaptic circuit
- Different portions of subfields showed different patterns of FC with neocortex

## Introduction

The hippocampus has been implicated in supporting multiple cognitive functions including episodic memory (Scoville and Milner, 1957), the imagination of fictitious and future experiences (Hassabis et al., 2007; Addis et al., 2007), spatial navigation (Chersi and Burgess, 2015; Maguire et al., 2006), visual perception (Lee et al., 2012; McCormick et al., 2017; Mullally et al., 2012) mind-wandering (McCormick et al., 2018; Smallwood et al., 2016; Karapanagiotidis et al., 2016) and decision making (McCormick et al., 2016; Mullally et al., 2014). The hippocampus is a heterogenous structure comprising multiple subregions including the dentate gyrus (DG), cornu ammonis (CA) 1-4, prosubiculum, subiculum, presubiculum, parasubiculum and the uncus. We currently lack a detailed understanding of how different parts of the human hippocampus interact with each other and with neighbouring cortical regions to support key functions such as memory.

The primary input to the hippocampus is via the entorhinal cortex (ENT), the source of the canonical tri-synaptic pathway. The ENT primarily innervates the DG and, from here, intra-hippocampal connectivity is generally acknowledged to follow a unidirectional pathway through the CA regions to the subiculum, the primary region of efferent projection from the hippocampus (Duvernoy et al., 2013; Aggleton and Christiansen, 2015). While this canonical circuitry is not in question, anatomical evidence has shown that extra-hippocampal regions including the ENT, perirhinal (PRC), posterior parahippocampal (PHC) and retrosplenial (RSC) cortices interact directly with specific hippocampal subfields, bypassing the canonical hippocampal pathway in both rodents (Agster and Burwell, 2013) and non-human primates (Witter and Amaral, 1991; Leonard et al., 1995; Aggleton, 2012; Kobayashi and Amaral, 2007). Moreover, tract tracing studies in non-human primates have revealed intra-subfield gradients of connectivity along the anterior-posterior axis of the hippocampus (Insausti and Munoz, 2001). This suggests that different portions of hippocampal subfields may preferentially interact with other brain regions. This is not surprising, given the known genetic, anatomical and functional differentiations along the long axis of the hippocampus (see Fanselow and Dong, 2009; Poppenck et al., 2013; Strange et al., 2014 for reviews).

One way to investigate homologues of these anatomical frameworks in vivo in the human brain is to characterise patterns of functional connectivity (FC) using resting state functional magnetic resonance imaging (rsfMRI). While rsfMRI FC often reflects anatomical pathways, its statistical dependencies are not limited to the underlying anatomy (Honey et al., 2009; 2010). Thus, rsfMRI FC has the additional benefit of reflecting potential functional relationships between brain regions. However, due largely to the technical difficulties inherent to investigating human hippocampal subfields using MRI, only a small number of studies have investigated rsfMRI FC of hippocampal subfields, and there has been no systematic examination of FC of the different portions of each subfield along the anterior-posterior axis of the human hippocampus.

Of the extant rsfMRI studies that considered human hippocampal subfield FC in some shape or form, several focussed on intra-medial temporal lobe (MTL) FC, including hippocampal subfields. They observed that activity was highly correlated between hippocampal subfields, and that functional networks within the MTL may divide into separate ‘functional modules’, one broadly consisting of hippocampal subfields and another consisting of parahippocampal regions (Shah et al., 2017; Lacy and Stark, 2012; although this is not a perfect dichotomy, see Shah et al., 2017). Other studies have predominantly focused on identifying hippocampal subfield FC changes as predictors of cognitive decline in age-related disorders (de Flores et al., 2017; Wang et al., 2015; Bai et al., 2011).

There have been some reports that patterns of FC for each hippocampal subfield broadly reflect those observed in studies of whole hippocampus network connectivity (Wang et al., 2015; Bai et al., 2011). By contrast, other studies have observed differences in patterns of hippocampal subfield FC. For example, de Flores et al. (2017) found that a combined DG/CA4/3/2 region of interest (ROI) was preferentially connected with left anterior cingulate, temporal and occipital regions. CA1 was preferentially connected with amygdala and occipital regions, and the subiculum (encompassing the subiculum, presubiculum and parasubiculum) was preferentially connected with angular gyrus, precuneus, putamen, posterior cingulate and frontal regions. Concordant with these findings, Vos de Wael et al. (2018) observed that subiculum activity was correlated with activity in regions associated with the default mode network, activity in a combined CA1-3 ROI was more strongly correlated with that of somatosensory and limbic regions, while the DG/CA4 showed patterns of connectivity with few cortical regions.

Of the very small number of studies examining FC in different parts of a subfield, it has been reported that the anterior portions of human subiculum and CA1 showed preferential FC with PRC while posterior portions exhibited preferential FC with PHC (Maas et al., 2015; Libby et al., 2012). Vos de Wael et al. (2018) observed complex variations in FC along the anterior-posterior axis of hippocampal subfields, but did not describe in detail exactly which cortical regions were associated with different portions along each hippocampal subfield gradient.

In the current study we aimed to systematically characterise not only the FC of each hippocampal subfield in its entirety (i.e. the total extent of the subfield along the longitudinal axis) but also the FC of different portions of each subfield along the anterior-posterior axis of the human hippocampus. We used a combination of high resolution structural MR imaging and high resolution rsfMRI to examine FC between the subfields themselves – DG/CA4, CA3/2, CA1, subiculum, pre- and parasubiculum, and the uncus – and subfield FC with key neighbouring regions in the memory/navigation network, specifically, ENT, PRC, PHC and also the RSC, due to its well documented anatomical connectivity with specific hippocampal subregions, in particular, the subiculum, pre- and parasubiculum (Dalton et al., 2017; Aggleton and Christiansen, 2015; Kravitz et al., 2011).

Of note, previous studies of hippocampal subfield FC have generally incorporated the entire subicular complex into a single subiculum mask (inclusive of the subiculum, presubiculum and parasubiculum; Shah et al., 2017; de Flores et al., 2017). However, different portions of the subicular complex show different patterns of anatomical (Kobayashi and Amaral, 2007; van Groen and Wyss, 1992, 2003) and functional (Maas et al., 2015) connectivity in humans and non-humans. Of particular interest, an anterior medial portion of the subicular complex encompassing the pre- and parasubiculum (hereafter referred to as the pre/parasubiculum), may facilitate elements of visual scene processing (Dalton and Maguire, 2017; Zeidman et al., 2015a,b) putatively through direct anatomical and functional connectivity with the RSC (Dalton and Maguire, 2017; Kravitz et al., 2011; Kobayashi and Amaral, 2007).

Anatomical data from non-human primates suggest that the RSC is a key node of the parieto-medial temporal visuospatial processing pathway and projects directly to the pre/parasubiculum (Kravitz et al., 2011). However, evidence for a functional link between the pre/parasubiculum and RSC in the human brain is lacking. Investigation of a functional homologue for this documented anatomical link was, therefore, of particular interest in the current study. With these considerations in mind, we utilised recently developed guidelines to differentiate the lateral subiculum ‘proper’ from the more medially located pre/parasubiculum (Dalton et al., 2017) and so, for the first time, separately investigated FC of the subiculum and pre/parasubiculum.

We also investigated for the first time FC of the uncus, a complex and understudied portion of the human hippocampus. From an anatomical perspective, the uncus also contains subfields, but these show modifications in cyto- and chemo-architecture compared to ‘typical’ hippocampal subfields (Ding and van Hoesen, 2015; McLardy, 1963) leading some anatomists to differentiate hippocampal subfields which lie in the uncus – uncal subfields – from those that lie in the ‘typical’ hippocampus (Ding and Van Hoesen, 2015). In addition, regions of the uncus also express differentiable patterns of anatomical connectivity in the primate brain (Insausti and Munoz, 2001; Rosene and van Hoesen, 1987).

Considering our hypotheses first in terms of FC between the whole subfields themselves, previous studies have generally observed activity between hippocampal subfields to be highly correlated. Here, we predicted the strength of correlations in activity between subfields would largely reflect our current understanding of the canonical intra-hippocampal circuitry (Duvernoy et al., 2013). Consequently, we hypothesised that: activity in DG/CA4 would most strongly correlate with activity in CA3/2; CA3/2 with DG/CA4 and CA1; CA1 with CA3/2 and subiculum; subiculum with CA1 and pre/parasubiculum; and pre/parasubiculum with subiculum.

In terms of FC of the whole hippocampal subfields with our cortical ROIs, we had specific hypotheses based primarily on patterns of anatomical connectivity reported in the non-human primate and rodent literatures. We predicted that: activity in DG/CA4 would correlate with activity in ENT (Duvernoy et al., 2013); CA3/2 activity would not correlate with activity in extra-hippocampal ROIs; activity in CA1 and subiculum would more strongly correlate with activity in ENT, PRC and PHC (Aggleton and Christiansen, 2015; Aggleton, 2012; Agster and Burwell, 2013; Rosene and van Hoesen, 1977; Kravitz et al., 2011); and in light of the rationale outlined above, we predicted that pre/parasubiculum activity would correlate with RSC, PHC (Kravitz et al., 2011; Kobayashi and Amaral, 2007; Van Groen and Wyss, 1992, 2003; Insausti and Munoz., 2001; Morris et al., 1999) and ENT activity (Huang et al., 2017; van Groen and Wyss, 1990; Caballero-Bleda and Witter, 1993; van Haeften 1997; Honda, 2004, 2011). Many of these anatomical connections were recently identified in the human brain by Zeineh et al., (2017) who conducted a study utilising polarised light imaging to visualise white matter pathways in post mortem human tissue. We did not have specific hypotheses relating to the FC of the uncus because, from both anatomical connectivity and functional perspectives, this region is understudied. Analyses relating to the uncus were, therefore, exploratory in nature.

Considering intra-subfield FC, to our knowledge, no study has investigated FC between different portions of the same hippocampal subfield. Beaujoin et al. (2018) conducted a multimodal analysis of hippocampal subfield anatomical connectivity in post mortem human hippocampus with ultra-high field MRI (11.7 Tesla). They reported that patterns of anatomical connectivity within hippocampal subfields extend along the longitudinal axis of the hippocampus. After splitting the hippocampus into three portions (head, body and tail) they found that adjacent portions of each subfield were anatomically connected (i.e. head with body; body with tail) but distant portions were not (i.e. head with tail). It is currently unknown whether there are functional homologues for these patterns of intra-subfield anatomical connectivity in the human brain. Taking these observations into account, we predicted that within each subfield, adjacent portions would be functionally correlated while more distant portions would not.

To our knowledge, no study has investigated FC between portions of each hippocampal subfield and different portions of other subfields along the longitudinal axis of the hippocampus. Observations from the non-human primate anatomical literature show that projections from hippocampal subfields extend bi-directionally along the longitudinal axis of the hippocampus (Kondo et al., 2008). Supporting this observation in humans, Beaujoin et al.’s (2018) structural MRI work revealed connections that extend through the length of the hippocampus, but that different portions of each hippocampal subfield had stronger anatomical connectivity with associated subfields in the same portion of the hippocampus and less with associated subfields in more distant portions of the hippocampus (i.e. DG/CA4 in the hippocampal head was correlated with CA3/2 in the hippocampal head but not with CA3/2 in the more distant body or tail portions of the hippocampus). Based on this recently-characterised anatomical framework, we predicted that different portions of each hippocampal subfield would have stronger anatomical connectivity with other subfields in the same or adjacent portions of hippocampus and not with subfields in more distant portions.

We also had specific predictions concerning FC of different portions of each subfield down the long axis of the hippocampus with the cortical ROIs. Based on the few extant investigations of hippocampal subfield long axis FC in humans (Vos de Wael et al., 2018; Maas et al., 2015; Libby et al., 2012) and the non-human primate anatomical literature noted above, we hypothesised that: activity in anterior portions of DG/CA4, CA1, subiculum and pre/parasubiculum would be correlated with ENT activity; the activity in anterior portions of CA1 and subiculum would be correlated with activity in PRC; activity in posterior portions of CA1 and subiculum would be correlated with PHC activity; and activity in anterior and posterior portions of the pre/parasubiculum would be correlated with activity in RSC and PHC.

To summarise, in this study we aimed to provide a comprehensive description of FC between the human hippocampal subregions themselves, and also their FC with neighbouring cortical regions. We did this at the whole subfield level and also for different portions of each subfield down the hippocampal long axis. Other novel aspects of this human hippocampal FC study included our separation of DG/CA4 from CA3/2, separation of the pre/parasubiculum from the subiculum, and the inclusion of the uncus as a separate ROI.

## Methods

### Participants

Twenty healthy, right handed participants took part in the study (11 females, mean age 23 years, SD 3.4). All gave written informed consent to participate in accordance with the University College London research ethics committee.

### Data acquisition and preprocessing

Structural and functional MRI data were acquired using a 3T Siemens Trio scanner (Siemens, Erlangen, Germany) with a 32-channel head coil within a partial volume centred on the temporal lobe that included the entire extent of the temporal lobes and our other ROIs.

Structural images were collected using a single-slab 3D T2-weighted turbo spin echo sequence with variable flip angles (SPACE) (Mugler et al., 2000) in combination with parallel imaging, to simultaneously achieve a high image resolution of ~500 μm, high sampling efficiency and short scan time while maintaining a sufficient signal-to-noise ratio (SNR). After excitation of a single axial slab the image was read out with the following parameters: resolution = 0.52 × 0.52 × 0.5 mm, matrix = 384 × 328, partitions = 104, partition thickness = 0.5 mm, partition oversampling = 15.4%, field of view = 200 × 171 mm 2, TE = 353 ms, TR = 3200 ms, GRAPPA × 2 in phase-encoding (PE) direction, bandwidth = 434 Hz/pixel, echo spacing = 4.98 ms, turbo factor in PE direction = 177, echo train duration = 881, averages = 1.9, plane of acquisition = sagittal. For reduction of signal bias due to, for example, spatial variation in coil sensitivity profiles, the images were normalized using a prescan, and a weak intensity filter was applied as implemented by the scanner’s manufacturer. To improve the SNR of the anatomical image, three scans were acquired for each participant, coregistered and averaged. Additionally, a whole brain 3D FLASH structural scan was acquired with a resolution of 1 × 1 × 1 mm.

Functional data were acquired using a 3D echo planar imaging (EPI) sequence which has been demonstrated to yield improved BOLD sensitivity compared to 2D EPI acquisitions (Lutti et al., 2013). Image resolution was 1.5 × 1.5 × 1.5 mm and the field-of-view was 192mm in-plane. Images were acquired in the sagittal plane. Forty slices were acquired with 20% oversampling to avoid wraparound artefacts due to the imperfect slab excitation profile. The echo time (TE) was 37.30 ms and the volume repetition time (TR) was 3.65s. Parallel imaging with GRAPPA image reconstruction (Griswold et al., 2002) acceleration factor 2 along the phase-encoding direction was used to minimise image distortions and yield optimal BOLD sensitivity. The dummy volumes necessary to reach steady state and the GRAPPA reconstruction kernel were acquired prior to the acquisition of the image data as described in Lutti et al. (2013). Correction of the distortions in the EPI images was implemented using B0-field maps obtained from double-echo FLASH acquisitions (matrix size 64×64; 64 slices; spatial resolution 3 × 3 × 3 mm; short TE=10 ms; long TE=12.46 ms; TR=1020 ms) and processed using the FieldMap toolbox in SPM (Hutton et al., 2002). Two hundred and five volumes were acquired with the scan lasting just under 13 minutes.

Preprocessing of structural and rsfMRI data was conducted using SPM12 (www.fil.ion.ac.uk/spm). All images were first bias-corrected, to compensate for image inhomogeneity associated with the 32 channel head coil (van Leemput et al., 1999). Fieldmaps were collected and used to generate voxel displacement maps. EPIs were then realigned to the first image and unwarped using the voxel displacement maps calculated above. The three high-resolution structural images were averaged to reduce noise, and co-registered to the whole brain structural FLASH scan. EPIs were also co-registered to the whole brain structural scan. In order to keep the EPI signal within each hippocampal subfield mask as pure as possible, no spatial smoothing was applied for these analyses, and all analyses were conducted in native space.

### Segmentation of hippocampal subfields

For each participant, we first manually delineated hippocampal subfields, bilaterally, on native space high resolution structural images according to the methodology described by Dalton et al. (2017) using the ITK Snap software version 3.2.0 (Yushkevich et al., 2006). Masks were created for the following subregions: DG/CA4, CA3/2, CA1, subiculum, pre/parasubiculum and uncus. To assess intra-rater reliability, five hippocampi were re-segmented 6 months apart and showed high concordance between segmentations at the two time points. Comparisons between the two segmentations were conducted using the Dice overlap metric (Dice, 1945) to produce a score between 0 (no overlap) and 1 (perfect overlap). Intra-rater reliability was 0.86 for DG/CA4, 0.71 for CA3/2, 0.81 for CA1, 0.79 for subiculum, 0.72 for pre/parasubiculum and 0.83 for the uncus. These values are equivalent to those reported in the extant literature (e.g. Bonnici et al., 2012; Polombo et al., 2013).

The gradient nature of connectivity along hippocampal subfields is well documented in both the anatomical (Insausti and Munoz, 2001; Beaujoin et al., 2018) and functional (Vos de Wael et al., 2018; Maas et al., 2015; Libby et al., 2012) literatures (reviewed in Strange et al., 2014; Poppenk et al., 2013). However, we do not have clear demarcations from either anatomy or function to guide the investigation of hippocampal subfields along their long axis. Hence, an often-used method takes the final slice of the uncus as a demarcation point for sectioning the hippocampus into two broad portions, anterior and posterior hippocampus (Zeidman et al., 2015b; Poppenck et al., 2013). While anatomically useful, this demarcation method may be problematic from a functional perspective. We have consistently observed a functional cluster in the medial hippocampus which extends across this demarcation point in tasks relating to scene-based cognition (Dalton et al., 2018; Zeidman et al., 2015a,b; Zeidman and Maguire, 2016). Hence, we believe that this portion of the hippocampus may represent a functional module which, when utilising the uncus-based anatomical demarcation point, would potentially be split between two separate ROIs. We, therefore, developed a novel method of demarcation which, while not designed to capture the gradient nature of connectivity, did allow us to sample broad portions of each subfield while ensuring this region was kept intact (Fig. 1).

**Fig. 1.**
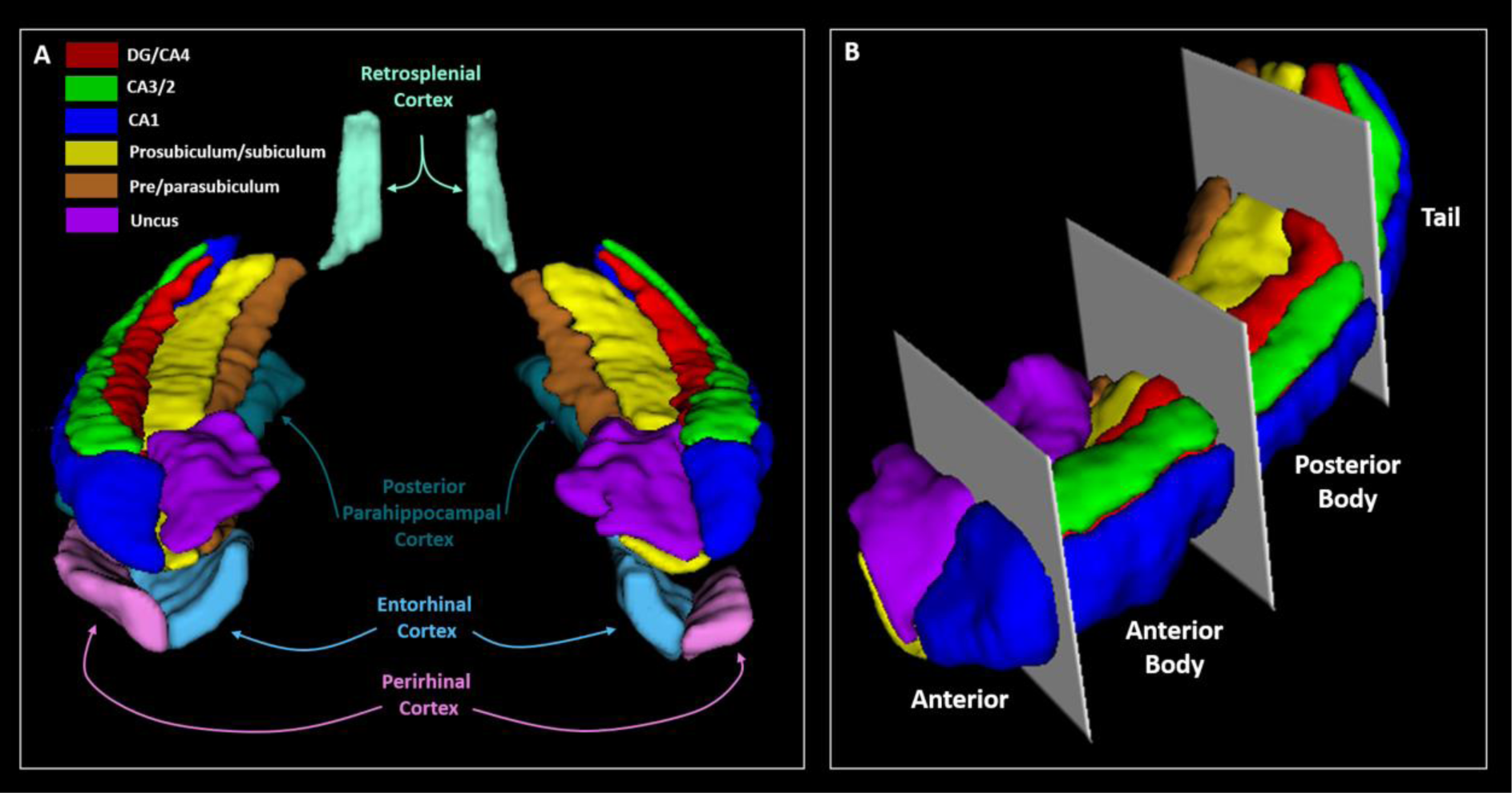
Regions of interest. **A.** A representative 3D model, viewed from an anterior perspective, of our primary regions of interest. **B.** A 3D representation, viewed from an antero-lateral perspective, of the boundaries we used to create anterior, anterior body, posterior body and tail portions of each hippocampal subfield.

We divided each subfield either into 4 (for CA1, subiculum and pre/parasubiculum), into 3 (for DG/CA4 and CA3/2) or into 2 (for the uncus) separate sections along its longitudinal axis (anterior (A), anterior body (AB), posterior body (PB) and tail (T); see Fig. 1B). For the anterior masks, the anterior boundary was the first slice of the hippocampus and the posterior boundary was the slice preceding the first slice of the dentate gyrus. This resulted in a mean of 15.5 (SD 3.3) slices in the anterior mask. The tail mask encompassed the posterior most 15 slices of the hippocampus. We had initially planned to use the crus of the fornix as the anterior demarcation for the tail masks but found that, due to individual variability in hippocampal morphology and flexure of the posterior hippocampus, this resulted in some participants having very few slices within the tail mask. In order to ensure that the tail mask contained an equivalent number of slices across participants we set the anterior most slice of the posterior portion to 15 slices anterior to and including the final slice of the hippocampus. The remaining slices were summed and split in half to create the anterior body and posterior body masks. This resulted in a mean of 22.9 (SD 2.4) and 22.5 (SD 2.4) slices in the anterior body and posterior body respectively. The average number of functional voxels contained within each subfield portion are provided in Table 1. Figure 2 shows the precise alignment between our structural and functional scans.

**Fig. 2.**
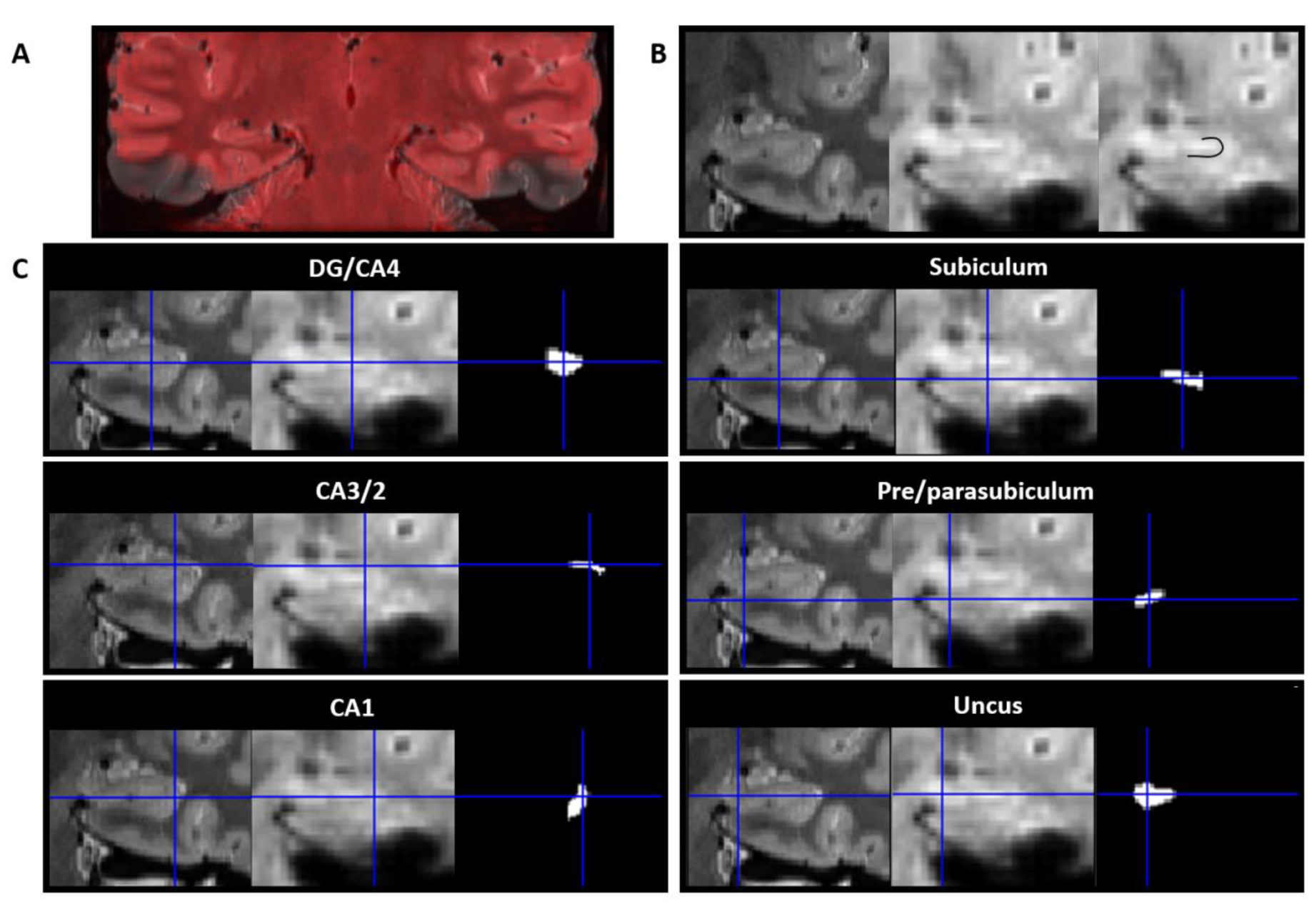
Alignment of the structural and functional scans with the hippocampal subfield ROIs. All images are presented in the coronal plane. **A.** Representative example of alignment between the high resolution partial volume T2 structural (greyscale) and functional (red) scans. Areas of signal dropout in the inferior lateral temporal lobes are clearly visible. **B.** Representative examples of the structural (left; 0.5mm isotropic voxels) and functional (middle; 1.5mm isotropic voxels) scans focussed on the hippocampus. Of note, the SLRM (comprised of the stratum radiatum and stratum lacunosum moleculare) is clearly visible on both scans (right; the same functional image as shown in middle panel with the SLRM overlaid with black line to aid visualisation). **C.** Representative examples of alignment between the high resolution structural scan (left), functional scan (middle) and functional ROI (right) for each hippocampal subfield.

**Table 1.**
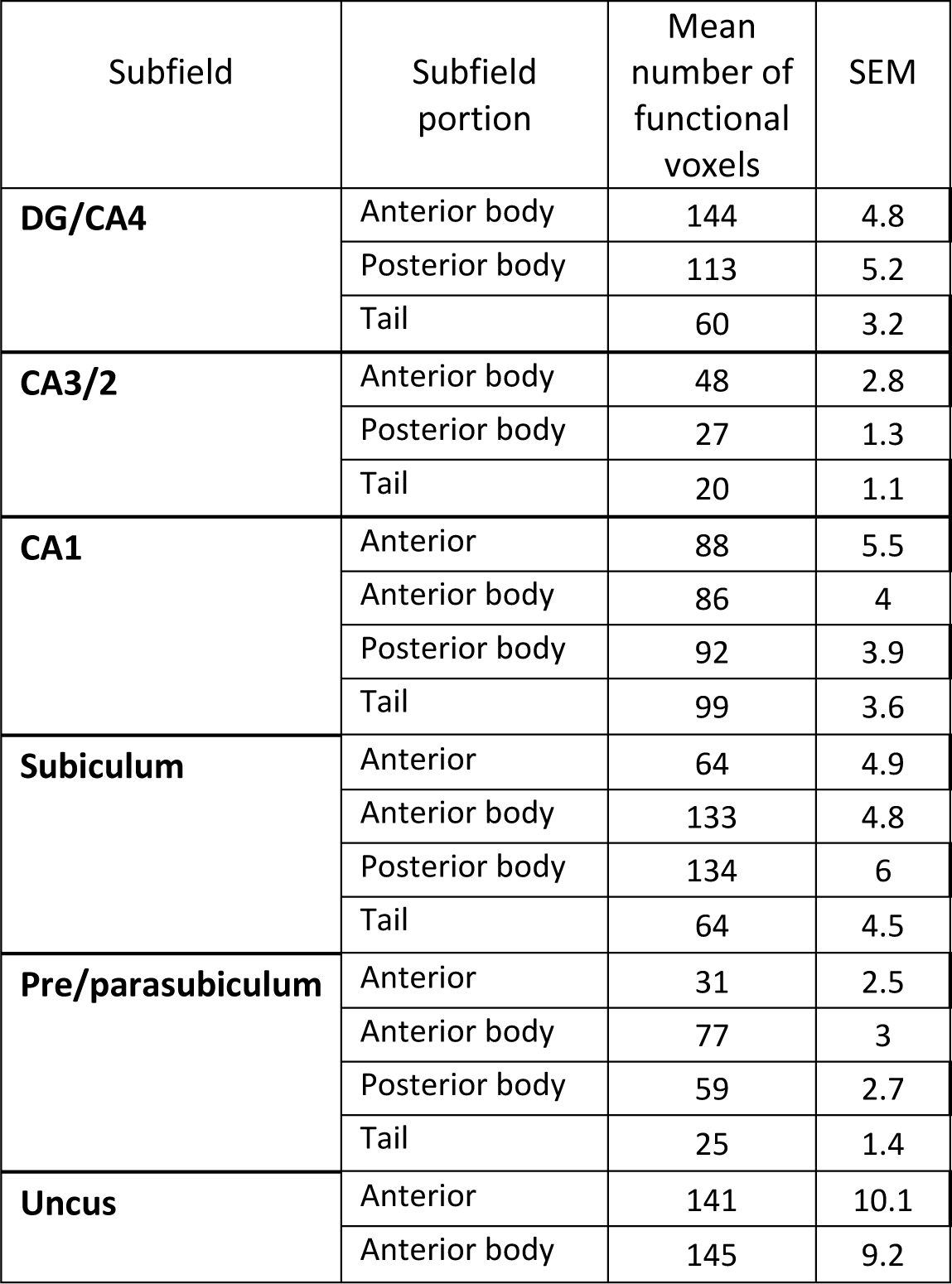
Mean number of functional voxels in each hippocampal ROI.

| Subfield | Subfield portion | Mean number of functional voxels | SEM |
| --- | --- | --- | --- |
| **DG/CA4** | Anterior body | 144 | 4.8 |
|  | Posterior body | 113 | 5.2 |
|  | Tail | 60 | 3.2 |
| **CA3/2** | Anterior body | 48 | 2.8 |
|  | Posterior body | 27 | 1.3 |
|  | Tail | 20 | 1.1 |
| **CA1** | Anterior | 88 | 5.5 |
|  | Anterior body | 86 | 4 |
|  | Posterior body | 92 | 3.9 |
|  | Tail | 99 | 3.6 |
| **Subiculum** | Anterior | 64 | 4.9 |
|  | Anterior body | 133 | 4.8 |
|  | Posterior body | 134 | 6 |
|  | Tail | 64 | 4.5 |
| **Pre/parasubiculum** | Anterior | 31 | 2.5 |
|  | Anterior body | 77 | 3 |
|  | Posterior body | 59 | 2.7 |
|  | Tail | 25 | 1.4 |
| **Uncus** | Anterior | 141 | 10.1 |
|  | Anterior body | 145 | 9.2 |

### Segmentation of extra-hippocampal ROIs

The ENT, PRC and PHC were segmented using the guidelines laid out by Augustinack et al. (2013), Fischl et al. (2009) and Berron et al. (2017). The anterior portions of ENT and PRC were generally prone to signal dropout on the fMRI scans. We, therefore, only included in our analyses portions of these subfields which clearly lay posterior to areas of signal dropout. We return to this point in the Discussion. To our knowledge, no guide to segmenting the human RSC on MRI scans is available. Consequently, we used the cytological investigation of the human RSC by Vogt et al. (2001) and the Allen Brain Atlas http://atlas.brain-map.org to gain insights into the likely location of the RSC. The RSC mask reflected our best attempt to remain as faithful as possible to the histological investigations of RSC. It was restricted to the thin strip of cortical tissue lying on the ventral bank of the cingulate gyrus and posterior to the corpus callosum, as described in detail by Vogt et al. (2001). Importantly, we did not include more posterior portions of the cingulate gyrus, and were careful not to include the callosal sulcus. Of note, recent evidence suggests that dorsal and ventral portions of the RSC may display distinct functional characteristics (Burles et al., 2018). In the current study, only ventral portions of the RSC were included owing to the partial volume.

### Data analysis

We used the CONN toolbox version 14 for rsfMRI data analysis http://www.nitrc.org/projects/conn. The data were temporally bandpass filtered (0.01 – 0.1 Hz) and corrected for white matter and ventricular signal. To create FC matrices, time series of voxels within each of the ROIs were averaged and correlated with the averaged time series of all other ROIs resulting in correlation coefficients which were then transformed using Fisher’s z calculation. Rather than using simple bivariate correlations, we used semi-partial correlations which allowed us to identify the ‘unique’ contribution of a given source on a target area. Of note, semi-partial correlations are computed between unmodified and other residualised variables, essentially regressing out or controlling contributions of other additional variables. Therefore, for each seed analysis in turn, slightly different values were regressed out, resulting in test statistics that vary marginally in their magnitude. That is, the semi-partial correlations between source region A and target region B might be slightly different than the semi-partial correlation between source region B and target region A. The resulting semi-partial ROI-to-ROI correlation matrices from the native space first-level analyses were further averaged at the second level in order to examine group effects. Importantly, this ROI-to-ROI approach allowed us to test hypotheses regarding FC between each ROI and all other ROIs using minimally preprocessed data (i.e. unsmoothed and not normalised). This approach minimised the mixing of BOLD signal between adjacent subfields. Any functional voxels overlapping an ROI border were assigned to whichever ROI contained the majority of its volume. Inevitably, there were differences between subfields, and portions of subfields, in terms of the number of functional voxels, given their size differences. What relationship this has, if any, to the number of functional connections identified or the strength of those connections is currently unknown. This issue should be explored in future studies.

First, we conducted ‘whole subfield’ analyses with 10 bilateral ROIs (DG/CA4, CA3/2, CA1, subiculum, pre/parasubiculum, uncus, ENT, PRC, PHC and RSC).

We then conducted ‘longitudinal axis’ analyses with 24 bilateral ROIs ([AB, PB and T DG/CA4], [AB, PB and T CA3/2], [A, AB, PB and T CA1], [A, AB, PB and T subiculum], [A, AB, PB and T pre/parasubiculum], [A and AB uncus], ENT, PRC, PHC and RSC). For both sets of analyses, ROI-to-ROI results were corrected for multiple comparisons and reported when significant at a level of p < 0.05 false discovery rate (FDR) corrected (Chumbley et al., 2010).

## Results

### Whole subfield analyses

We first analysed the FC of each hippocampal subfield in its entirety, in terms of FC with other subfields and with the cortical ROIs. The results are summarised in Figure 3 and Table 2, which also includes the results of the statistical analyses.

**Fig. 3.**
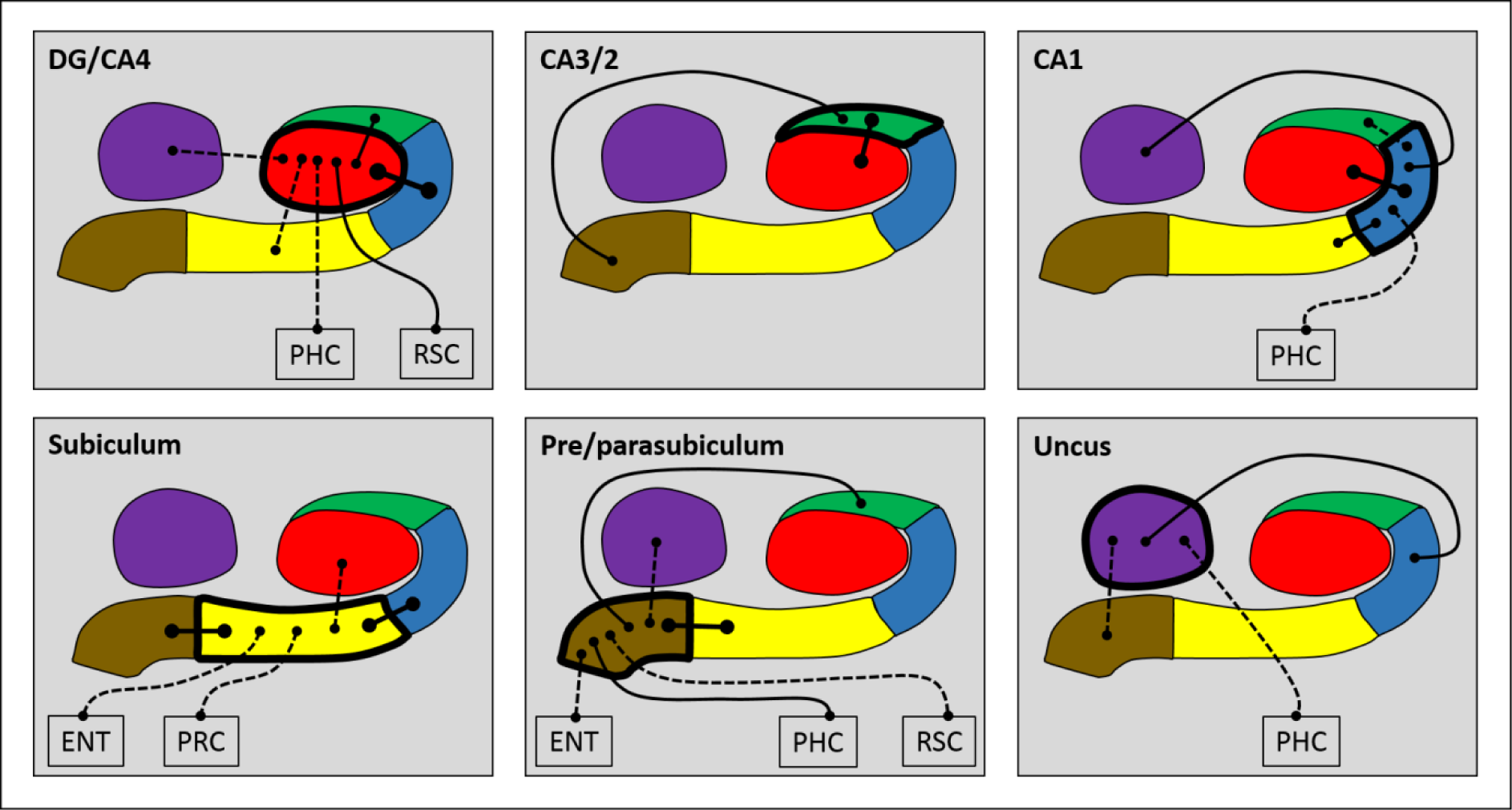
Results of the whole subfield analyses. The relevant subfield in each panel is outlined in a thick black line. The black lines with circular termini represent significant correlations of activity with the activity in other hippocampal subfields and/or extra-hippocampal ROIs at p < 0.05 FDR corrected. Connection strength is indicated by line type. Thick unbroken lines = t > 10; thin unbroken lines = t > 5; thin broken lines = t < 5. DG/CA4 (red), CA3/2 (green), CA1 (blue), subiculum (yellow), pre/parasubiculum (brown), uncus (purple); ENT=entorhinal cortex, PRC=perirhinal cortex, PHC=posterior parahippocampal cortex, RSC=retrosplenial cortex.

**Table 2.** Results of the whole subfield analyses.

| Seed ROI | Significant Target ROI's | T - Statistic | p – FDR corrected |
| --- | --- | --- | --- |
| **DG/CA4** | CA3/2 | T (19) = 8.24 | < 0.0001 |
|  | CA1 | T (19) = 12.56 | < 0.0001 |
|  | Subiculum | T (19) = 4.69 | 0.0004 |
|  | Uncus | T (19) = 2.33 | 0.0397 |
|  | Perirhinal cortex | T (19) = -2.95 | 0.0149 |
|  | Parahippocampal cortex | T (19) = 2.38 | 0.0397 |
|  | Retrosplenial cortex | T (19) = 5.39 | < 0.0001 |
| **CA3/2** | DG/CA4 | T (19) = 8.90 | < 0.0001 |
|  | Pre/parasubiculum | T (19) = 5.46 | < 0.0001 |
|  | Uncus | T (19) = -5.37 | < 0.0001 |
|  | Retrosplenial cortex | T (19) = -2.61 | 0.0386 |
| **CA1** | DG/CA4 | T (19) = 11.49 | < 0.0001 |
|  | CA3/2 | T (19) = 2.52 | 0.0270 |
|  | Subiculum | T (19) = 9.13 | < 0.0001 |
|  | Pre/parasubiculum | T (19) = -4.92 | 0.0002 |
|  | Uncus | T (19) = 6.93 | < 0.0001 |
|  | Entorhinal cortex | T (19) = -4.08 | 0.0012 |
|  | Parahippocampal cortex | T (19) = 2.52 | 0.0270 |
| **Subiculum** | DG/CA4 | T (19) = 4.77 | 0.0004 |
|  | CA1 | T (19) = 10.16 | < 0.0001 |
|  | Pre/parasubiculum | T (19) = 11.78 | < 0.0001 |
|  | Entorhinal cortex | T (19) = 2.55 | 0.0349 |
|  | Perirhinal cortex | T (19) = 2.66 | 0.0349 |
| **Pre/parasubiculum** | CA3/2 | T (19) = 5.48 | < 0.0001 |
|  | CA1 | T (19) = -4.98 | 0.0002 |
|  | Subiculum | T (19) = 11.42 | 0.0000 |
|  | Uncus | T (19) = 4.13 | 0.0009 |
|  | Entorhinal cortex | T (19) = 3.72 | 0.0019 |
|  | Parahippocampal cortex | T (19) = 5.74 | < 0.0001 |
|  | Retrosplenial cortex | T (19) = 4.40 | 0.0006 |
| **Uncus** | CA3/2 | T (19) = -5.82 | < 0.0001 |
|  | CA1 | T (19) = 6.37 | < 0.0001 |
|  | Pre/parasubiculum | T (19) = 4.69 | 0.0005 |
|  | Parahippocampal cortex | T (19) = 3.50 | 0.0053 |
|  | Retrosplenial cortex | T (19) = -2.96 | 0.0143 |

Negative correlations are shown in italics.

**<u>DG/CA4</u>** was significantly correlated with CA3/2, CA1, subiculum, uncus, PHC and RSC.

**<u>CA3/2</u>** was correlated with DG/CA4 and the pre/parasubiculum.

**<u>CA1</u>** was correlated with DG/CA4, subiculum, uncus, CA3/2 and PHC.

**<u>Subiculum</u>** was correlated with pre/parasubiculum, CA1, DG/CA4, ENT and PRC.

**<u>Pre/parasubiculum</u>** was correlated with the subiculum, CA3/2, uncus, ENT, PHC and RSC.

The **<u>uncus</u>** was correlated with CA1, pre/parasubiculum and PHC.

These results suggest that each hippocampal subfield had a unique pattern of FC with other hippocampal subfields, and that each subfield showed a different pattern of FC with the cortical ROIs.

### Longitudinal axis analyses

Next, we analysed FC for different portions of each hippocampal subfield along its anterior-posterior axis, in terms of FC between different portions of a subfield, FC with other subfield portions and with the cortical ROIs. These results are summarised in Figures 4-9 and Table 3, which also includes the results of the statistical analyses.

**Fig. 4.**
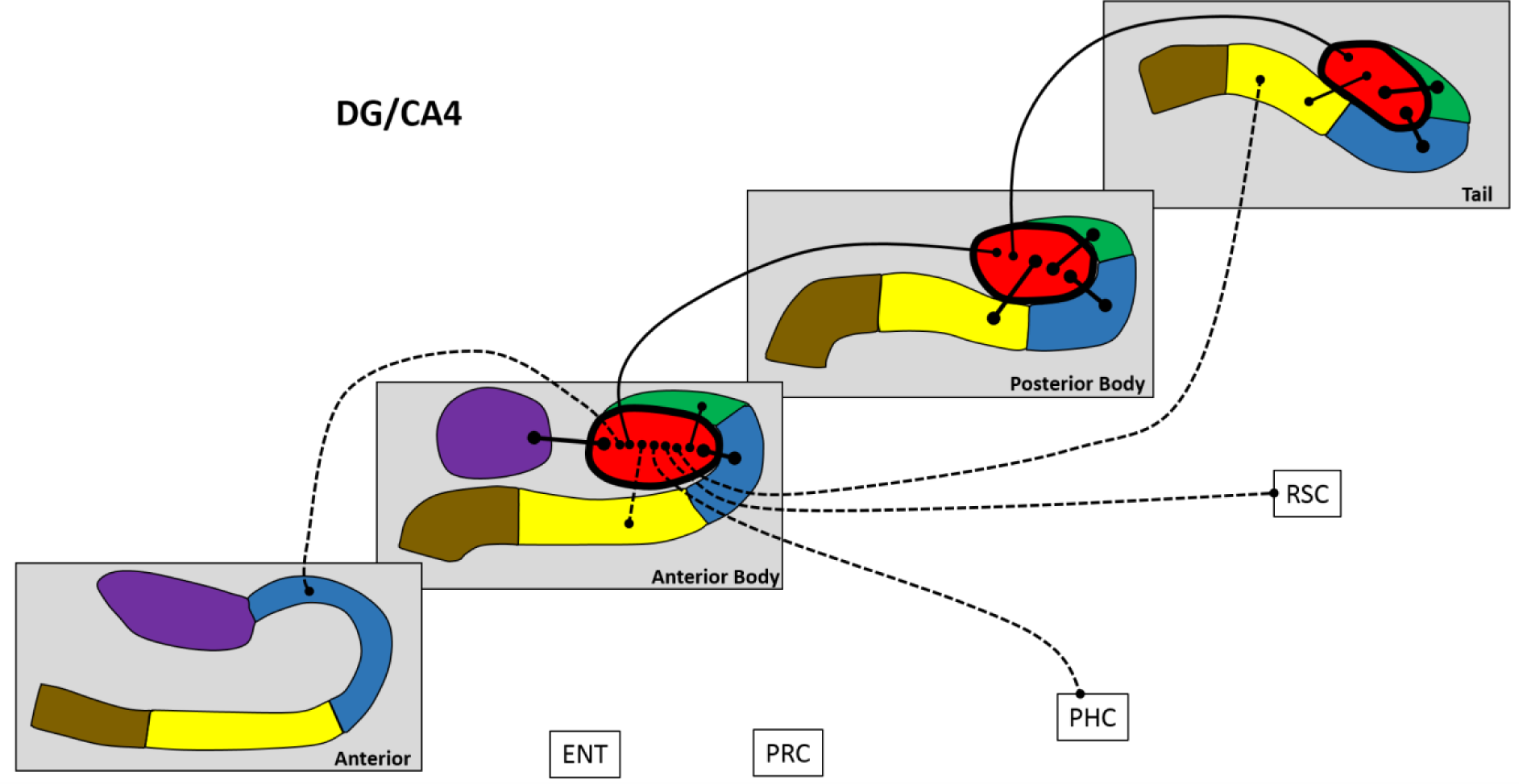
Results of longitudinal axis analysis for DG/CA4. The relevant subfield in each panel is outlined in a thick black line. The black lines with circular termini represent significant correlations of activity at p < 0.05 FDR. Connection strength is indicated by line type. Thick unbroken lines = t > 10; thin unbroken lines = t > 5; thin broken lines = t < 5. DG/CA4 (red), CA3/2 (green), CA1 (blue), subiculum (yellow), pre/parasubiculum (brown), uncus (purple); ENT=entorhinal cortex, PRC=perirhinal cortex, PHC=posterior parahippocampal cortex, RSC=retrosplenial cortex.

**Fig. 5.**
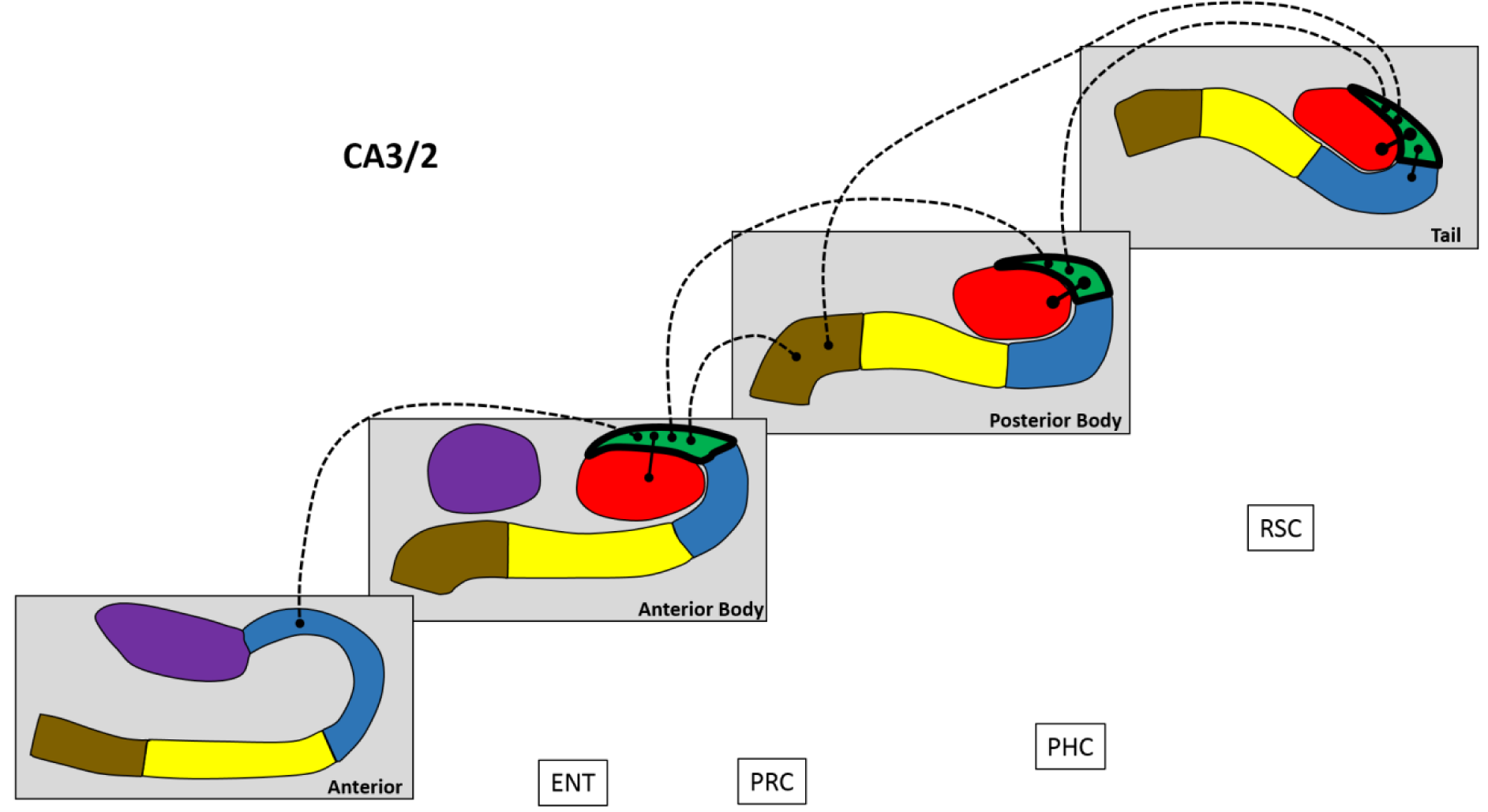
Results of longitudinal axis analysis for CA3/2. The relevant subfield in each panel is outlined in a thick black line. The black lines with circular termini represent significant correlations of activity at p < 0.05 FDR. Connection strength is indicated by line type. Thick unbroken lines = t > 10; thin unbroken lines = t > 5; thin broken lines = t < 5. DG/CA4 (red), CA3/2 (green), CA1 (blue), subiculum (yellow), pre/parasubiculum (brown), uncus (purple); ENT=entorhinal cortex, PRC=perirhinal cortex, PHC=posterior parahippocampal cortex, RSC=retrosplenial cortex.

**Fig. 6.**
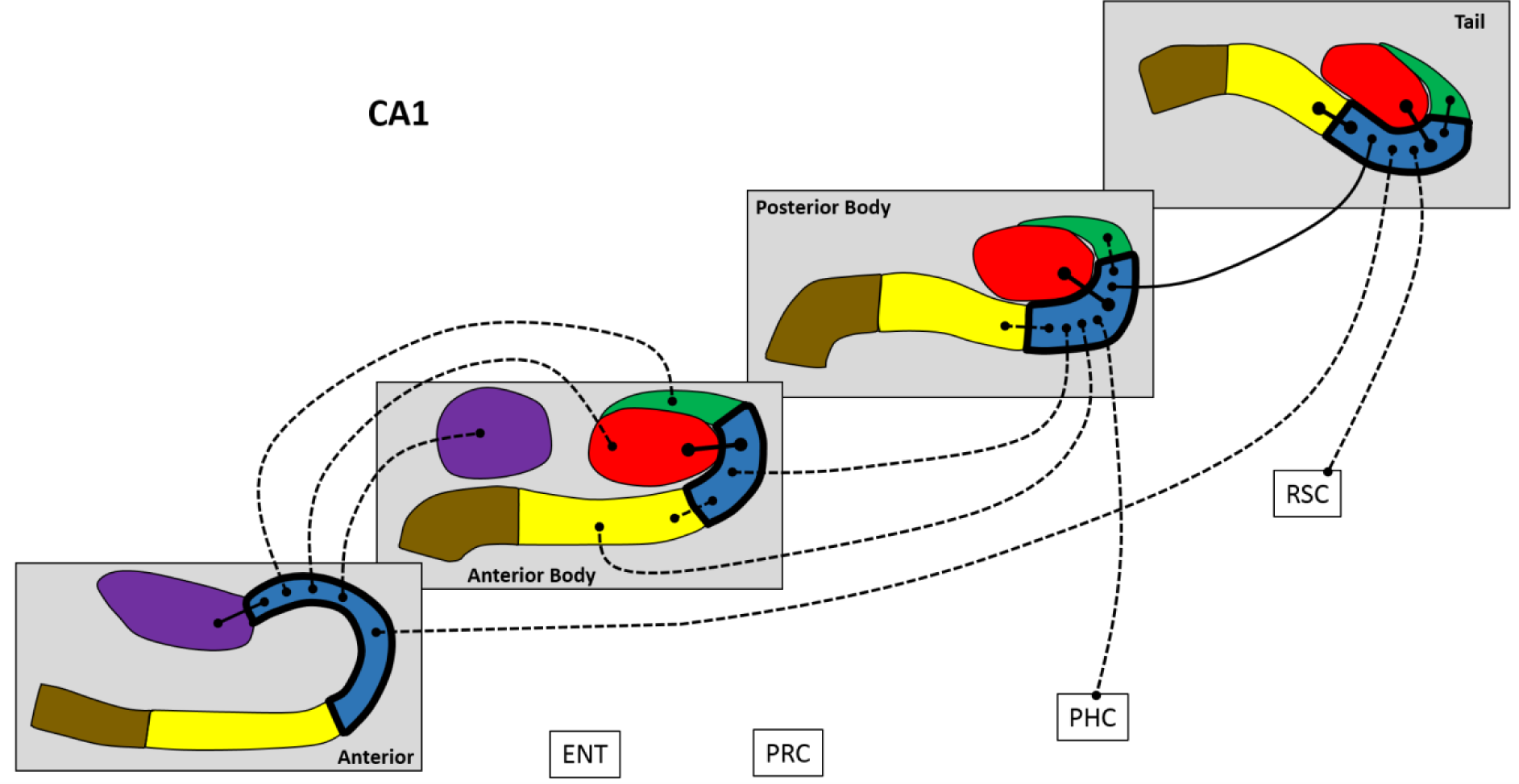
Results of longitudinal axis analysis for CA1. The relevant subfield in each panel is outlined in a thick black line. The black lines with circular termini represent significant correlations of activity at p < 0.05 FDR. Connection strength is indicated by line type. Thick unbroken lines = t > 10; thin unbroken lines = t > 5; thin broken lines = t < 5. DG/CA4 (red), CA3/2 (green), CA1 (blue), subiculum (yellow), pre/parasubiculum (brown), uncus (purple); ENT=entorhinal cortex, PRC=perirhinal cortex, PHC=posterior parahippocampal cortex, RSC=retrosplenial cortex.

**Fig. 7.**
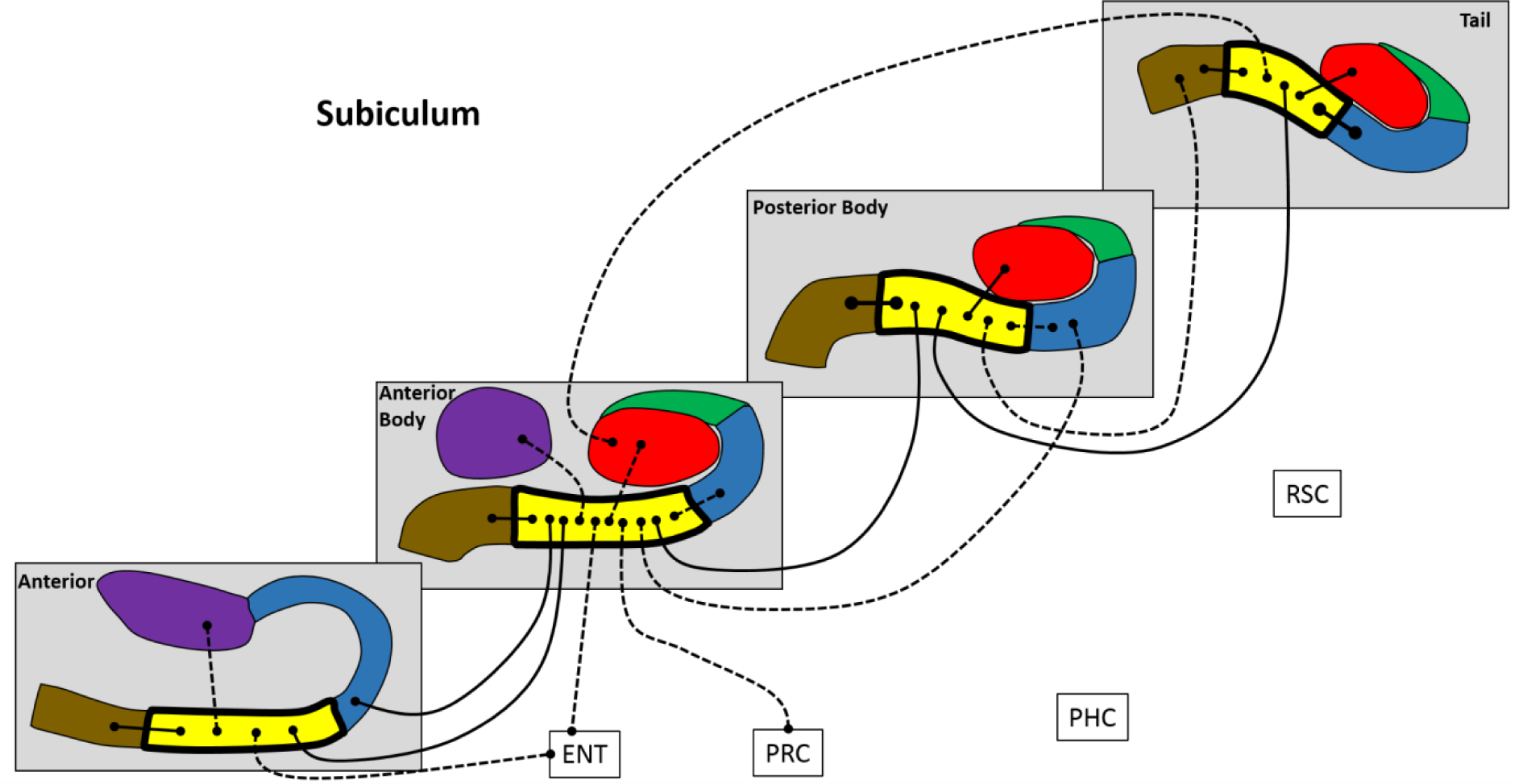
Results of longitudinal axis analysis for the subiculum. The relevant subfield in each panel is outlined in a thick black line. The black lines with circular termini represent significant correlations of activity at p < 0.05 FDR. Connection strength is indicated by line type. Thick unbroken lines = t > 10; thin unbroken lines = t > 5; thin broken lines = t < 5. DG/CA4 (red), CA3/2 (green), CA1 (blue), subiculum (yellow), pre/parasubiculum (brown), uncus (purple); ENT=entorhinal cortex, PRC=perirhinal cortex, PHC=posterior parahippocampal cortex, RSC=retrosplenial cortex.

**Fig. 8.**
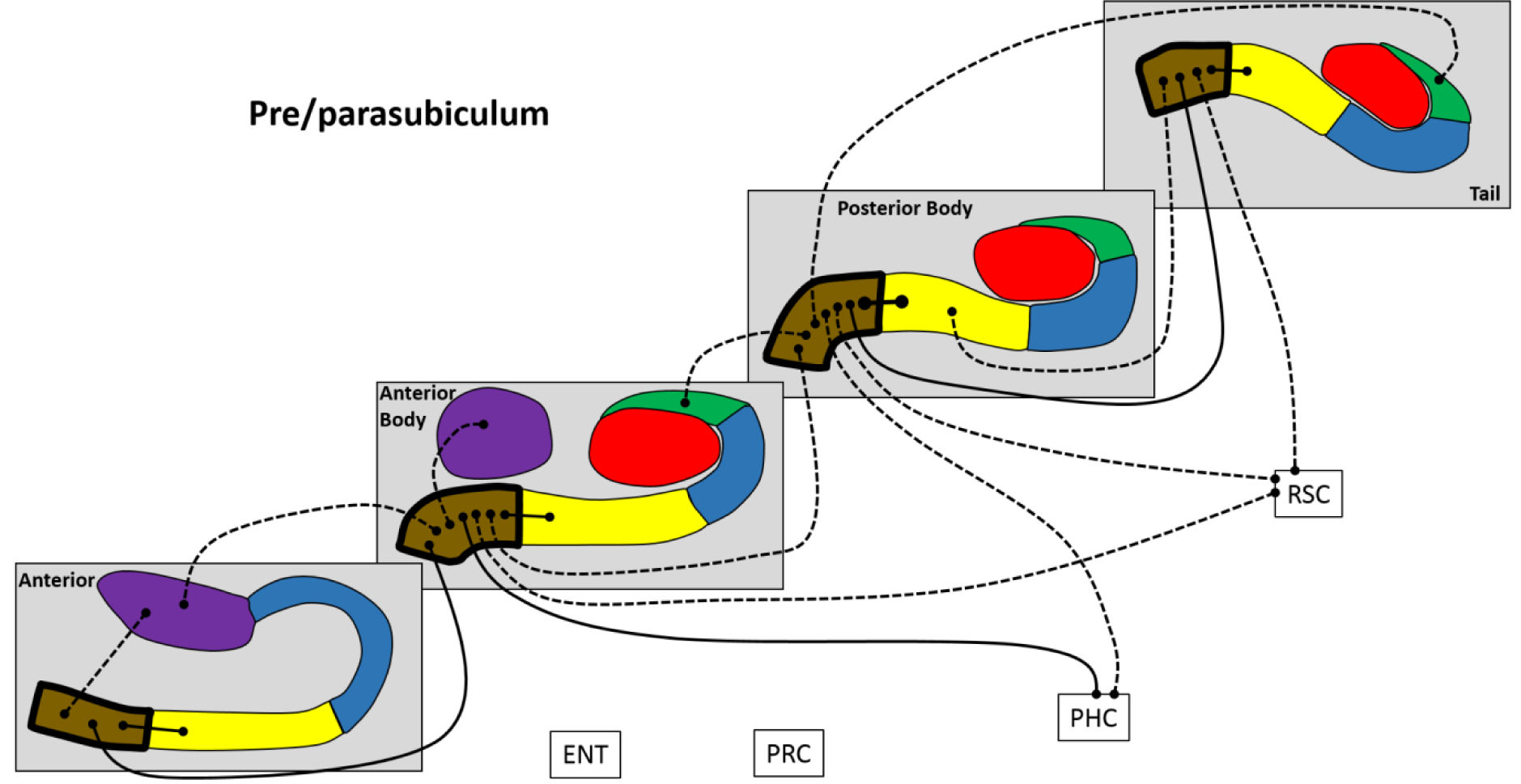
Results of longitudinal axis analysis for the pre/parasubiculum. The relevant subfield in each panel is outlined in a thick black line. The black lines with circular termini represent significant correlations of activity at p < 0.05 FDR. Connection strength is indicated by line type. Thick unbroken lines = t > 10; thin unbroken lines = t > 5; thin broken lines = t < 5. DG/CA4 (red), CA3/2 (green), CA1 (blue), subiculum (yellow), pre/parasubiculum (brown), uncus (purple); ENT=entorhinal cortex, PRC=perirhinal cortex, PHC=posterior parahippocampal cortex, RSC=retrosplenial cortex.

**Fig. 9.**
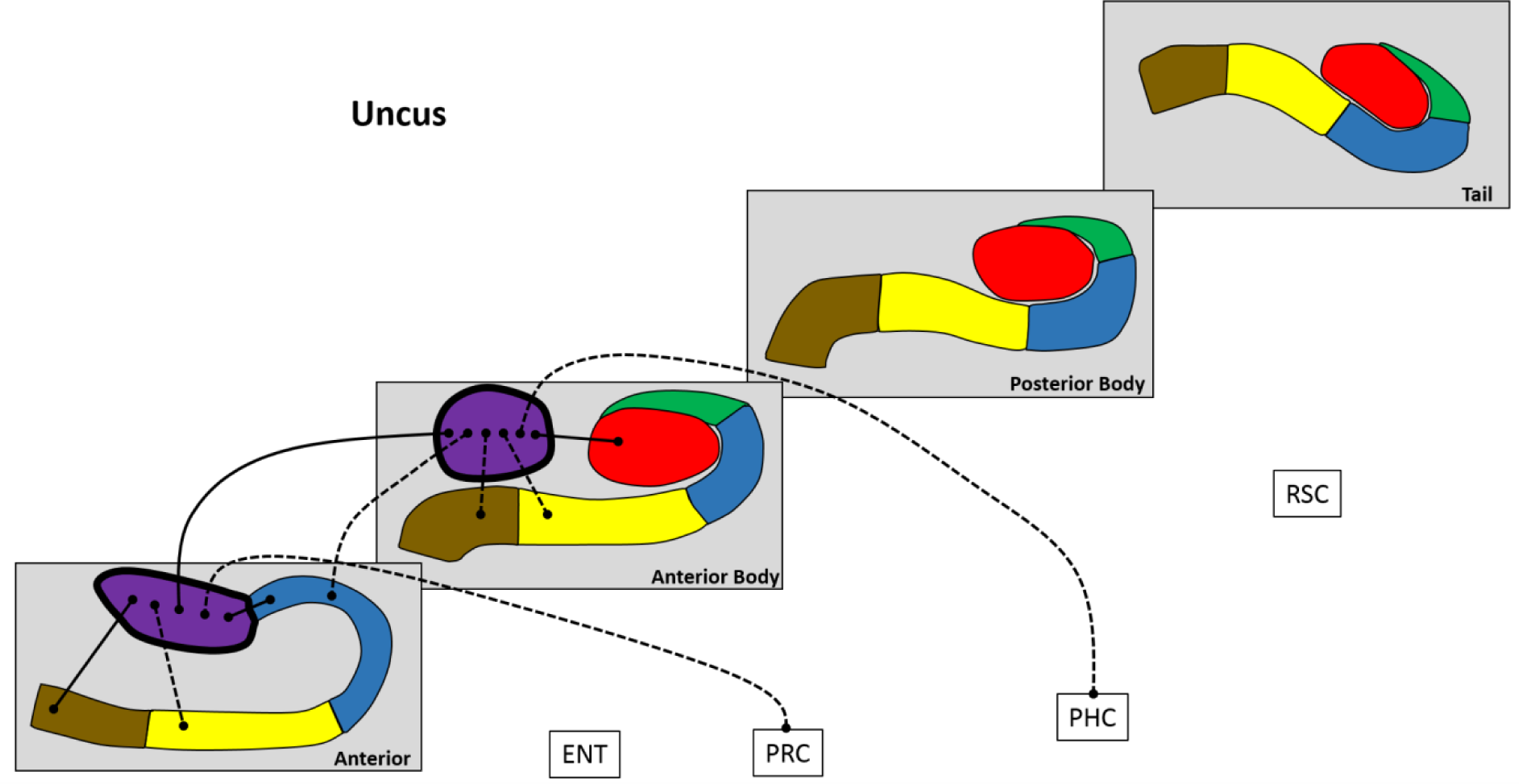
Results of longitudinal axis analysis for the uncus. The relevant subfield in each panel is outlined in a thick black line. The black lines with circular termini represent significant correlations of activity at p < 0.05 FDR. Connection strength is indicated by line type. Thick unbroken lines = t > 10; thin unbroken lines = t > 5; thin broken lines = t < 5. DG/CA4 (red), CA3/2 (green), CA1 (blue), subiculum (yellow), pre/parasubiculum (brown), uncus (purple); ENT=entorhinal cortex, PRC=perirhinal cortex, PHC=posterior parahippocampal cortex, RSC=retrosplenial cortex.

**Table 3.** Results of the longitudinal axis analyses.

| Seed ROI | Significant Target ROI's | T - statistic | P – FDR corrected |
| --- | --- | --- | --- |
| **DG/CA4** |  |  |  |
| Anterior body | Anterior CA1 | T (19) = 3.50 | 0.0079 |
|  | Anterior subiculum | T (19) = -2.48 | 0.0473 |
|  | Anterior body CA3/2 | T (19) = 5.89 | < 0.0001 |
|  | Anterior body CA1 | T (19) = 10.65 | < 0.0001 |
|  | Anterior body subiculum | T (19) = 2.60 | 0.0405 |
|  | Anterior body uncus | T (19) = 11.96 | < 0.0001 |
|  | Posterior body DG/CA4 | T (19) = 8.36 | < 0.0001 |
|  | Posterior body CA3/2 | T (19) = -5.52 | < 0.0001 |
|  | Tail subiculum | T (19) = 3.53 | 0.0079 |
|  | Parahippocampal cortex | T (19) = 3.02 | 0.0191 |
|  | Retrosplenial cortex | T (19) = 2.99 | 0.0191 |
| Posterior body | Anterior body DG/CA4 | T (19) = 8.88 | < 0.0001 |
|  | Posterior body CA3/2 | T (19) = 12.73 | < 0.0001 |
|  | Posterior body CA1 | T (19) = 13.89 | < 0.0001 |
|  | Posterior body subiculum | T (19) = 10.18 | < 0.0001 |
|  | Tail DG/CA4 | T (19) = 9.37 | < 0.0001 |
| Tail | Posterior body DG/CA4 | T (19) = 8.58 | < 0.0001 |
|  | Tail CA3/2 | T (19) = 11.41 | < 0.0001 |
|  | Tail CA1 | T (19) = 11.52 | < 0.0001 |
|  | Tail subiculum | T (19) = 7.42 | < 0.0001 |
| **CA3/2** |  |  |  |
| Anterior body | Anterior CA1 | T (19) = 3.13 | 0.0211 |
|  | Anterior uncus | T (19) = -3.62 | 0.0085 |
|  | Anterior body DG/CA4 | T (19) = 5.39 | 0.0008 |
|  | Posterior body CA3/2 | T (19) = 4.87 | 0.0012 |
|  | Posterior body CA1 | T (19) = -4.62 | 0.0014 |
|  | Posterior body pre/parasubiculum | T (19) = 3.85 | 0.0062 |
| Posterior body | Anterior body DG/CA4 | T (19) = -4.64 | 0.0013 |
|  | Anterior body CA3/2 | T (19) = 4.90 | 0.0012 |
|  | Posterior body DG/CA4 | T (19) = 11.43 | < 0.0001 |
|  | Tail CA3/2 | T (19) = 4.55 | 0.0013 |
| Tail | Posterior body CA3/2 | T (19) = 5.07 | 0.0005 |
|  | Posterior body pre/parasubiculum | T (19) = 3.19 | 0.0279 |
|  | Tail DG/CA4 | T (19) = 10.86 | < 0.0001 |
|  | Tail CA1 | T (19) = 7.49 | < 0.0001 |
| **CA1** |  |  |  |
| Anterior | Anterior uncus | T (19) = 5.26 | < 0.0001 |
|  | Anterior body DG/CA4 | T (19) = 3.38 | < 0.0001 |
|  | Anterior body CA3/2 | T (19) = 2.96 | < 0.0001 |
|  | Anterior body uncus | T (19) = 3.29 | < 0.0001 |
|  | Tail CA1 | T (19) = 3.91 | < 0.0001 |
|  | Entorhinal cortex | T (19) = -3.85 | 0.0004 |
| Anterior body | Anterior body DG/CA4 | T (19) = 10.11 | < 0.0001 |
|  | Anterior body subiculum | T (19) = 4.78 | 0.0015 |
|  | Posterior body CA1 | T (19) = 4.30 | 0.0030 |
|  | Posterior body pre/parasubiculum | T (19) = -3.04 | 0.0391 |
| Posterior body | Anterior body CA3/2 | T (19) = -4.22 | 0.0025 |
|  | Anterior body CA1 | T (19) = 4.15 | 0.0025 |
|  | Anterior body subiculum | T (19) = 3.85 | 0.0042 |
|  | Anterior body pre/parasubiculum | T (19) = -3.38 | 0.0094 |
|  | Posterior body DG/CA4 | T (19) = 13.42 | < 0.0001 |
|  | Post body CA3/2 | T (19) = 2.66 | 0.0298 |
|  | Posterior body subiculum | T (19) = 4.40 | 0.0024 |
|  | Posterior body pre/parasubiculum | T (19) = -3.09 | 0.0152 |
|  | Tail CA3/2 | T (19) = -2.73 | 0.0276 |
|  | Tail CA1 | T (19) = 5.35 | 0.0004 |
|  | Tail pre/parasubiculum | T (19) = -3.36 | 0.0094 |
|  | Parahippocampal cortex | T (19) = 2.84 | 0.0240 |
| Tail | Anterior CA1 | T (19) = 3.69 | 0.0072 |
|  | Anterior body pre/parasubiculum | T (19) = -2.85 | 0.0394 |
|  | Posterior body CA1 | T (19) = 6.08 | < 0.0001 |
|  | Tail DG/CA4 | T (19) = 11.69 | < 0.0001 |
|  | Tail CA3/2 | T (19) = 7.48 | < 0.0001 |
|  | Tail subiculum | T (19) = 10.65 | < 0.0001 |
|  | Retrosplenial cortex | T (19) = 2.71 | 0.0456 |
| **Subiculum** |  |  |  |
| Anterior | Anterior pre/parasubiculum | T (19) = 7.23 | < 0.0001 |
|  | Anterior uncus | T (19) = 3.95 | < 0.0001 |
|  | Anterior body subiculum | T (19) = 7.40 | < 0.0001 |
|  | Entorhinal cortex | T (19) = 3.05 | 0.0383 |
| Anterior body | Anterior CA1 | T (19) = 2.64 | 0.0443 |
|  | Anterior subiculum | T (19) = 7.31 | < 0.0001 |
|  | Anterior body DG/CA4 | T (19) = 2.60 | 0.0443 |
|  | Anterior body CA1 | T (19) = 5.35 | 0.0003 |
|  | Anterior body pre/parasubiculum | T (19) = 6.20 | < 0.0001 |
|  | Anterior body uncus | T (19) = 4.13 | 0.0026 |
|  | Posterior body CA1 | T (19) = 3.80 | 0.0046 |
|  | Posterior body subiculum | T (19) = 5.15 | 0.0003 |
|  | Entorhinal cortex | T (19) = 3.41 | 0.0095 |
|  | Perirhinal cortex | T (19) = 2.56 | 0.0443 |
| Posterior body | Anterior body subiculum | T (19) = 5.13 | 0.0003 |
|  | Anterior body uncus | T (19) = -4.60 | 0.0009 |
|  | Posterior body DG/CA4 | T (19) = 8.95 | < 0.0001 |
|  | Posterior body CA1 | T (19) = 4.24 | 0.0015 |
|  | Posterior body pre/parasubiculum | T (19) = 11.59 | < 0.0001 |
|  | Tail subiculum | T (19) = 7.95 | < 0.0001 |
|  | Tail pre/parasubiculum | T (19) = 4.43 | 0.0011 |
| Tail | Anterior body DG/CA4 | T (19) = 3.74 | 0.0064 |
|  | Posterior body subiculum | T (19) = 8.64 | < 0.0001 |
|  | Tail DG/CA4 | T (19) = 7.10 | < 0.0001 |
|  | Tail CA1 | T (19) = 10.49 | < 0.0001 |
|  | Tail pre/parasubiculum | T (19) = 8.52 | < 0.0001 |
| **Pre/parasubiculum** |  |  |  |
| **Anterior** | Anterior subiculum | T (19) = 7.22 | < 0.0001 |
|  | Anterior body pre/parasubiculum | T (19) = 5.23 | 0.0006 |
|  | Anterior uncus | T (19) = 4.85 | 0.0008 |
| Anterior body | Anterior pre/parasubiculum | T (19) = 5.16 | 0.0004 |
|  | Anterior uncus | T (19) = 2.56 | 0.0491 |
|  | Anterior body subiculum | T (19) = 6.16 | < 0.0001 |
|  | Anterior body uncus | T (19) = 3.70 | 0.0070 |
|  | Posterior body CA1 | T (19) = -3.33 | 0.0116 |
|  | Posterior body pre/parasubiculum | T (19) = 3.84 | 0.0064 |
|  | Tail CA1 | T (19) = -2.78 | 0.0346 |
|  | Parahippocampal cortex | T (19) = 6.16 | < 0.0001 |
|  | Retrosplenial cortex | T (19) = 3.41 | 0.0114 |
| Posterior body | Anterior body CA3/2 | T (19) = 3.89 | 0.0056 |
|  | Anterior body pre/parasubiculum | T (19) = 3.77 | 0.0059 |
|  | Posterior body CA1 | T (19) = -2.77 | 0.0349 |
|  | Posterior body subiculum | T (19) = 11.37 | < 0.0001 |
|  | Tail CA3/2 | T (19) = 3.04 | 0.0259 |
|  | Tail pre/parasubiculum | T (19) = 5.91 | < 0.0001 |
|  | Parahippocampal cortex | T (19) = 4.23 | 0.0035 |
|  | Retrosplenial cortex | T (19) = 2.92 | 0.0286 |
| Tail | Posterior body CA1 | T (19) = -3.44 | 0.0126 |
|  | Posterior body subiculum | T (19) = 4.46 | 0.0021 |
|  | Posterior body pre/parasubiculum | T (19) = 6.00 | < 0.0001 |
|  | Tail subiculum | T (19) = 8.20 | < 0.0001 |
|  | Retrosplenial cortex | T (19) = 3.51 | 0.0126 |
| **Uncus** |  |  |  |
| Anterior | Anterior CA1 | T (19) = 5.10 | 0.0005 |
|  | Anterior subiculum | T (19) = 3.88 | 0.0052 |
|  | Anterior pre/parasubiculum | T (19) = 5.26 | 0.0005 |
|  | Anterior body CA3/2 | T (19) = -3.83 | 0.0052 |
|  | Anterior body uncus | T (19) = 6.64 | < 0.0001 |
|  | Tail subiculum | T (19) = -2.85 | 0.0339 |
|  | Perirhinal cortex | T (19) = 2.99 | 0.0286 |
| Anterior body | Anterior CA1 | T (19) = 3.56 | 0.0068 |
|  | Anterior uncus | T (19) = 6.32 | < 0.0001 |
|  | Anterior body DG/CA4 | T (19) = 9.97 | < 0.0001 |
|  | Anterior body subiculum | T (19) = 4.21 | 0.0027 |
|  | Anterior body pre/parasubiculum | T (19) = 3.99 | 0.0032 |
|  | Posterior body subiculum | T (19) = -4.86 | 0.0008 |
|  | Parahippocampal cortex | T (19) = 3.96 | 0.0032 |

Negative correlations are shown in italics.

**<u>DG/CA4</u>** (Fig. 4):

Activity in the AB portion was significantly correlated with activity in A CA1, AB CA3/2, AB CA1, AB subiculum, AB uncus, PB DG/CA4, T subiculum, PHC and RSC.

The PB portion was associated with AB DG/CA4, PB CA3/2, PB CA1, PB subiculum, T DG/CA4.

The T portion was associated with PB DG/CA4, T CA3/2, T CA1 and T subiculum.

To summarise, in relation to FC between different portions of DG/CA4, each part was correlated with adjacent but not distant portions (e.g. the AB portion correlated with the PB portion but not the T portion). Considering FC with other subfields, the DG/CA4 in each portion of the hippocampus correlated with CA3/2, CA1 and subiculum within the same portion of the hippocampus, but rarely with more distant portions of these subfields. AB DG/CA4 was the only portion of the DG/CA4 to correlate with more distant portions of other subfields, specifically the A CA1 and T subiculum. AB DG/CA4 also correlated with AB uncus. In addition, AB DG/CA4 was the only portion of the DG/CA4 to correlate with the cortical ROIs, specifically PHC and RSC. This suggests that AB DG/CA4 may have a broader pattern of both intra- and extra-hippocampal FC compared with more posterior portions.

**<u>CA3/2</u>** (Fig. 5):

Activity in the AB portion was correlated with A CA1, AB DG/CA4, PB CA3/2 and PB pre/parasubiculum.

The PB portion was associated with AB CA3/2, PB DG/CA4 and T CA3/2.

The T portion was correlated with PB CA3/2, PB pre/parasubiculum, T DG/CA4 and T CA1.

To summarise, in relation to FC between different portions of CA3/2, each part was correlated with adjacent but not distant portions. Considering FC with other subfields, CA3/2 in each portion of the hippocampus correlated with DG/CA4 within the same portion of the hippocampus. The AB CA3/2 correlated with A CA1. The T CA3/2 correlated with the T CA1 and both the AB and T CA3/2 were correlated with the PB pre/parasubiculum. We found no evidence for CA3/2 correlations with any of the cortical ROIs.

**<u>CA1</u>** (Fig. 6):

The A portion was associated with A uncus, AB DG/CA4, AB CA3/2, AB uncus and T CA1.

The AB portion was associated with AB DG/CA4, AB subiculum, PB CA1.

The PB portion was associated with AB CA1, AB subiculum, PB DG/CA4, PB CA3/2, PB subiculum, T CA1 and PHC.

The T portion was associated with A CA1, PB CA1, T DG/CA4, T CA3/2,T subiculum and RSC.

To summarise, in relation to FC between different portions of CA1, AB and PB portions of CA1 were correlated only with adjacent portions. A and T CA1 were correlated with each other. Considering FC with other subfields, CA1 in each portion of the hippocampus correlated with DG/CA4 and subiculum within the same portion of the hippocampus but rarely with distant portions of these subfields. The only parts of CA1 to show FC with the cortical ROIs were PB CA1 with PHC and T CA1 with RSC.

**<u>Subiculum</u>** (Fig. 7):

The A portion was associated with AB subiculum, A pre/parasubiculum, A uncus and ENT.

The AB portion was correlated with A CA1, A subiculum, AB DG/CA4, AB CA1, AB pre/parasubiculum, AB uncus, PB CA1, PB subiculum, ENT and PRC.

The PB portion was associated with AB subiculum, PB DG/CA4, PB CA1, PB pre/parasubiculum, T subiculum and T pre/parasubiculum.

The T portion was associated with AB DG/CA4, PB subiculum, T DG/CA4, T CA1 and T pre/parasubiculum.

To summarise, in relation to FC between different portions of the subiculum, each portion was correlated with adjacent but not distant portions. Considering FC with other subfields, the subiculum in each portion of the hippocampus correlated with DG/CA4, CA1 and pre/parasubiculum within the same portion of the hippocampus, but also often with distant portions of these subfields. In terms of FC with the cortical ROIs, A and AB subiculum correlated with ENT while AB subiculum correlated with PRC.

**<u>Pre/parasubiculum</u>** (Fig. 8):

The A portion was associated with A subiculum, A uncus and AB pre/parasubiculum.

The AB portion was correlated with A pre/parasubiculum, A uncus, AB subiculum, AB uncus, PB pre/parasubiculum, PHC and RSC.

The PB portion was associated with AB CA3/2, AB pre/parasubiculum, PB subiculum, T CA3/2, T pre/parasubiculum, PHC and RSC.

The T portion was associated with PB subiculum, PB pre/parasubiculum, T subiculum and RSC.

To summarise, in relation to FC between different portions of the pre/parasubiculum, each part was correlated with adjacent but not distant portions. Considering FC with other subfields, the pre/parasubiculum in each portion of the hippocampus correlated with subiculum only within the same portion of the hippocampus, with the exception of the T pre/parasubiculum which also correlated with PB subiculum. PB pre/parasubiculum correlated with AB and T portions of the CA3/2. The A pre/parasubiuculum correlated with A uncus while AB pre/parasubiculum correlated with both A and AB uncus. In terms of FC with the cortical ROIs, PB pre/parasubiculum correlated with PHC while AB, PB and T pre/parasubiculum correlated with RSC.

**<u>Uncus</u>** (Fig. 9):

The A portion was associated with A CA1, A subiculum, A pre/parasubiculum, AB uncus and PRC.

The AB portion was associated with A CA1, A uncus, AB DG/CA4, AB subiculum, AB pre/parasubiculum and PHC.

To summarise, in relation to FC between different portions of the uncus, the A and AB portions were correlated with each other. Considering FC with other subfields, both the A and AB uncus correlated with subiculum and pre/parasubiculum within the same portion of the hippocampus. Both A and AB uncus correlated with A CA1 while AB uncus correlated with AB DG/CA4. In terms of FC with the cortical ROIs, A uncus correlated with PRC while AB uncus correlated with PHC.

## Discussion

In this study we investigated the resting state FC of hippocampal subfields and specific extra-hippocampal ROIs by leveraging high resolution structural and functional MRI. We first probed the FC of each hippocampal subfield in its entirety (i.e. the total extent of each subfield along the longitudinal axis of the hippocampus). We examined how each subfield interacted with other hippocampal subfields and with neighbouring cortical areas. We then investigated the FC of the anterior, anterior body, posterior body and tail portions of each subfield. We aimed to characterise how each portion of each subfield interacted with other portions of the same subfield, with other portions of other subfields and, finally, with neighbouring cortical ROIs. Our results provide new insights into how human hippocampal subfields may functionally interact with each other along the anterior-posterior axis of the hippocampus during a resting state. In addition, we show that the ENT, PRC, PHC and RSC preferentially interact, not only with specific hippocampal subfields, but with specific portions of each subfield along the hippocampal longitudinal axis.

### Whole subfield analyses - intra-hippocampal FC

Previous studies investigating FC between hippocampal subfields have found them to be highly correlated with each other (Shah et al., 2017). In the current study, we used semi-partial correlations to identify the ‘unique’ contribution of a given source on a target area. In essence, this method represents the connectivity between two ROI’s after controlling for BOLD time series in all other ROI’s. Using this approach, we found that while hippocampal subfields were highly correlated with each other, when controlling for activity in all other subfields, each subfield also had a unique pattern of FC with other subfields.

For the most part, we observed functional homologues for the well characterised canonical intra-hippocampal anatomical circuitry (DG/CA4 → CA3/2 → CA1 → subiculum) in alignment with our predictions (e.g. DG/CA4 was correlated with CA3/2, CA1 was correlated with subiculum). However, in addition, we also documented some unexpected functional correlations. For instance, we observed an association between CA3/2 and pre/parasubiculum. A rationale for a functional association between these regions is unclear, and we are unaware of any previous reports of direct anatomical or functional interactions between the pre/parasubiculum and CA3/2 in the human brain. These observations suggest that FC between hippocampal subfields may extend beyond the current understanding of anatomical connectivity. We return to this point later.

### Whole subfield analyses - FC with cortical ROIs

In addition to having different patterns of FC with other subfields, each hippocampal subfield had a unique pattern of FC with the neighbouring ROIs. Largely aligning with our predictions, the ENT was associated with the subiculum and pre/parasubiculum; the PRC was associated with the subiculum; the PHC with the DG/CA4, CA1, pre/parasubiculum and uncus; and the RSC with the pre/parasubiculum and DG/CA4.

This is the first study to investigate FC of the pre/parasubiculum in the human brain. Concordant with one of our primary predictions, activity in the pre/parasubiculum was correlated with activity in RSC. The RSC is a key node of the parieto-medial temporal visuospatial processing pathway (Kravitz et al., 2011) and sends direct projections specifically to the pre/parasubiculum and CA1 hippocampal subfields and also to the PHC (Kobayashi and Amaral, 2007; Kravitz et al., 2011). While functional homologues for other components of the parieto-medial temporal visuospatial processing pathway have been identified in the human brain (Margulies et al., 2009; Caminiti et al., 2010; Rushworth et al., 2006), to our knowledge, this is the first identification of an explicit functional link between the human RSC and the pre/parasubiculum portion of the hippocampus. Our results support extant proposals that these regions may comprise an anatomical-functional unit (Barbas and Blatt, 1995; Golman-Rakic, 1984) related to elements of visuospatial processing (Dalton and Maguire., 2017; Kravitz et al., 2011).

We have previously outlined a potential rationale for the functional significance of the pre/parasubiculum in visuospatial processing in relation to scene-based cognition (Dalton and Maguire, 2017). In brief, we proposed that the pre/parasubiculum is a key hippocampal hub of the parieto-medial temporal visuospatial processing pathway and may be neuroanatomically determined to preferentially process integrated holistic representations of the environment. These representations, in turn, are held to underpin the scene-based cognition upon which episodic memory, imagination of fictitious or future scenarios and mind wandering are dependent (Maguire and Mullally, 2013; McCormick et al., 2018). The pre/parasubiculum may, therefore, play an important role in supporting these cognitive abilities. Our observation of a specific functional link between the RSC and pre/parasubiculum lends further support to this idea.

Previous studies of hippocampal FC do not differentiate between the pre/parasubiculum and subiculum ‘proper’ (Shah et al., 2017) and, therefore, necessarily refer to the entire ‘subicular complex’ as subiculum. Multiple elements of our results, however, clearly show that the pre/parasubiculum and subiculum proper have different patterns of FC. Together with the recent observation that even different regions of the pre/parasubiculum may facilitate distinct forms of mental imagery (Dalton et al., 2018), the current findings provide persuasive evidence that neuroimaging investigations of the hippocampus should, at the very least, differentiate between the pre/parasubiculum and the subiculum proper when investigating contributions of this complex region to human cognition. Indeed, Insausti et al. (2017) recently suggested that the concept of a ‘subicular cortex’ may be incorrect considering it combines the differentiable subicular allocortex (comprising 3 layers) and the pre/parasubicular periallocortex (comprising 6 layers). Our results provide support for this distinction at a functional level. While acknowledging that these regions can be difficult to differentiate on MRI, recent hippocampal segmentation protocols offer reliable methods for doing so (Dalton et al., 2017; Iglesias et al., 2015).

The interpretation of other interesting observations from the whole subfield analyses are facilitated by the results of the longitudinal axis analyses. We therefore discuss other elements of these results in the context of the longitudinal axis analyses below.

### Longitudinal axis analyses - intra-subfield FC

Considering the longitudinal axis analyses, we first investigated how each portion of each subfield interacted with other portions of the same subfield (e.g. FC between A, AB, PB and T portions of CA1). We observed that, within each subfield, not all portions were functionally correlated with each other, as would be expected if each subfield was functionally homogeneous. Rather, generally within each subfield, adjacent portions were correlated while distant portions were not. For example, AB DG/CA4 correlated with the adjacent PB DG/CA4 but not with the more distant T DG/CA4. This pattern was consistent for almost all subfields and implies a functional heterogeneity within each hippocampal subfield and that distant portions of each subfield may not be functionally ‘in synch’ with each other. These results mirror patterns of intra-subfield anatomical connectivity recently described by Beaujoin et al. (2018) who found that, when splitting the human hippocampus into three portions (head, body and tail), adjacent, but not distant, portions of each subfield were anatomically connected. Our FC results deviated from this pattern, however, in relation to CA1 where, in addition to the pattern described above, we observed an association between the more distant A and T portions. This observation is intriguing but difficult to interpret. Whether it suggests that CA1 plays a role in directly relaying information between the anterior and posterior hippocampus remains unclear. Fibres which run along the longitudinal axis of the human hippocampus have recently been directly observed in the human brain (Zeineh et al., 2017). These fibres may represent a mechanism by which distant portions of hippocampal subfields along their anterior-posterior axis could directly interact with each other. Further characterisation of these longitudinal fibres may elucidate whether a biological mechanism which facilitates direct communication between distant portions of hippocampal subfields exists or not. This CA1 finding also shows that it is unlikely our FC results were simply the result of proximity effects, considering the significant functional connectivity between these non-adjacent subregions (see more on this below). Moreover we also observed instances where directly-adjacent regions showed weak or no significant functional connectivity. Also of note, in order to minimise the mixing of BOLD signal between neighbouring subregions, we did not apply spatial smoothing to our functional data.

### Longitudinal axis analyses - intra-hippocampal interactions between subfields

We next investigated how each portion of each subfield interacted with different portions of other subfields. In accordance with our predictions, the functional homologues for the canonical anatomical framework observed in the whole subfield analysis were, for the most part, also observed within each portion of the hippocampus (e.g. DG/CA4 was correlated with CA3/2 within AB, PB and T portions of the hippocampus; see Figure 4). Beaujoin et al. (2018) have reported that different portions of human hippocampal subfields have stronger anatomical connectivity with associated subfields in the same portion of the hippocampus (e.g. strong anatomical connectivity between AB DG/CA4 and AB CA3/2) and less anatomical connectivity with associated subfields in more distant portions of the hippocampus (e.g. weak or no anatomical connectivity between AB DG/CA4 and T CA3/2). While our results broadly support this pattern at a functional level (see Figure 3), it is interesting to note that in addition to this pattern, FC between associated subfields was not invariably restricted to, and was sometimes not present within, the same portion of the hippocampus. For example, CA3/2 was correlated with CA1 within the tail of the hippocampus but not within the anterior body or posterior body of the hippocampus (see Figure 4). Rather, AB CA3/2 was correlated with the adjacent A CA1, and activity in T subiculum was correlated with AB DG/CA4. Indeed, compared to other subfields, the subiculum had more extensive patterns of FC with adjacent and distant portions of other subfields (see Figure 7) suggesting that this region may have a broader range of intra-hippocampal FC than other subfields. This is interesting considering the well documented role of the subiculum as the primary region of efferent projection from the hippocampus (Duvernoy et al., 2013; Aggleton and Christiansen, 2015). On a related note, Kondo et al. (2008) observed extensive versus limited interconnections in the posterior two-thirds versus the anterior one-third of the non-human primate hippocampus respectively (see Strange et al., 2014 for discussion). This pattern was not evident in the human brain either in the current study or in the data reported by Beaujoin et al., (2018). Whether this represents fundamental differences in patterns of intra-hippocampal connectivity between species or simply reflect methodological issues remains an open question.

Taken together, these observations suggest that while, for the most part, different portions of each hippocampal subfield have FC with associated subfields in the same portion of the hippocampus, specific portions of each subfield may have preferential FC with distant rather than nearby portions of associated subfields. From an anatomical perspective, such interactions may be facilitated by longitudinal fibres which project through the anterior-posterior axis of the hippocampus, supporting functional links between distant portions of hippocampal subfields (Zeineh et al., 2017; Beaujoin et al., 2018).

### Longitudinal axis analyses - hippocampal subfield FC with extra-hippocampal cortices

The longitudinal axis analyses offered additional insights into the nature of the FC patterns we identified at the whole subfield level in terms of FC with the neighbouring cortical ROIs. We found evidence that different portions of each subfield had different patterns of FC with these cortical regions. We discuss each of these observations in turn.

The results of our longitudinal axis analyses broadly support previous FC studies which suggested that the PRC and PHC disproportionately interact with anterior and posterior portions of the hippocampus respectively (Kahn et al., 2008), particularly in the subiculum and CA1 (Libby et al., 2012; Maas et al., 2015). Our findings extend this further by revealing that the AB portion of the subiculum and the A uncus were preferentially correlated with PRC. Interestingly, the anterior-most portion of the uncus (defined here as slices of the uncus which lie anterior to the emergence of the DG) is predominantly comprised of uncal subiculum and uncal CA1 (see Ding and van Hoesen, 2015). Our results are, therefore, in accordance with previous observations of an association between the PRC and anterior portions of subiculum and CA1 but, importantly, suggest that PRC may have preferential FC with uncal subiculum/CA1.

It is important to note that while we specifically predicted that PRC would be correlated with anterior portions of CA1 and subiculum, neuroanatomical evidence suggests a link between the PRC and the transition zone of the CA1 and subiculum, encompassing a region sometimes referred to as the prosubiculum. This CA1-subiculum transition area has been consistently observed in numerous tract tracing studies in the non-human primate and appears to have direct interactions with a number of brain regions including the PRC and cingulate cortex (Kondo et al., 2005; Insausti and Munoz, 2001; Vogt and Pandya, 1987), suggesting it may be a ‘hotspot’ of direct hippocampal connectivity. Taking into consideration the difficulties inherent in definitively identifying subfield borders on MRI, and that this hotspot lies at the transition between CA1 and subiculum, it is impossible to know whether this region lies within our subiculum or CA1 mask, or is split between the two masks. Future investigations of the FC of this CA1-subiculum transition area will be needed to explore this further.

As noted above, previous FC studies also suggest that PHC has preferential connectivity with posterior portions of CA1 and subiculum. Concordant with this, we observed a correlation between PB CA1 and PHC. In contrast, however, we did not observe an association between the subiculum and PHC. Rather, we noted that the AB and PB portions of the pre/parasubiculum were correlated with PHC. A reason for this discrepancy may lie in how the subiculum has been defined in previous studies where, as alluded to earlier, pre/parasubiculum is typically incorporated into a general subiculum region. By separating these areas, our data provide increased specificity on the nature of this functional interaction.

An interesting pattern was also observed in the A and AB portions of the uncus. The A uncus was associated with PRC while the more posteriorly located AB uncus was associated with PHC. To our knowledge, there are no previous studies of the FC of the human uncus and, therefore, this is the first report of a functional dissociation between its anterior and posterior portions. It must be noted that the uncus is a complex portion of the hippocampus which itself contains multiple subfields (Ding and Van Hoesen, 2015) that cannot be reliably differentiated on 3T MRI. Results relating to the uncus must, therefore, be interpreted with this in mind. The functional significance of the differences observed here are as yet unclear and further investigations of this complex region are clearly warranted.

Anterior portions of the subiculum were correlated with ENT, reflecting the well characterised anatomical connections between these regions (Aggleton, 2012; Aggleton and Christiansen, 2015). It was surprising, however, considering the documented associations between ENT and DG (Duvernoy et al., 2013), that we observed no significant correlation between the ENT and DG/CA4 in our analyses. Interestingly, a lack of FC between these regions has been noted in previous studies (e.g. Lacy and Stark, 2012) and could reflect methodological issues relating to fMRI signal dropout in anterior portions of the infero-temporal lobes. In the current study, we only included posterior portions of the ENT which showed no evidence of signal dropout within our ENT mask. However, considering the anatomical (Insausti et al., 2017) and functional (Maas et al., 2015) heterogeneity within the ENT, this method may not be adequate. Indeed, according to the framework recently proposed by Insausti et al. (2017), our ENT mask likely only included the ‘caudal’ and ‘caudal limiting’ portions of the ENT. Further advances in anatomical and functional parcellation (Maas et al., 2014, 2015) of the ENT may be key to future efforts to characterise how different functional components of this complex region interact with hippocampal subfields.

Finally, we observed correlations between the RSC and the AB, PB and T portions of the pre/parasubiculum and the T portion of the CA1. Anatomical frameworks of the parieto-medial temporal visuospatial processing pathway, described earlier, highlight that the RSC sends direct projections that specifically innervate the pre/parasubiculum and CA1 portions of the hippocampus (Dalton and Maguire, 2017; Kravitz et al., 2011). Our FC results dovetail with these documented patterns of anatomical connectivity and support this framework at a functional level.

It is important to note that the results of these longitudinal analysis show that correlations found at the whole subfield level may be driven by specific portions of a subfield rather than the whole. For example, the whole subfield analyses revealed that the pre/parasubiculum was correlated with CA3/2 while the longitudinal analysis revealed, more specifically, that the PB portion of the pre/parasubiculum was correlated with AB and T portions of CA3/2 while no other portion of the preparasubiculum was correlated with CA3/2. In a similar manner, while the whole subfield analyses revealed that the CA1 was correlated with PHC, the longitudinal analyses suggest that only the PB portion of CA1 was correlated with PHC. Patterns such as these underline the overarching theme of our results, namely that different portions of each subfield have different patterns of FC both with other hippocampal subfields and with neighbouring cortical ROI’s and that, where possible, this should be taken into account in future studies.

### Gradient nature of intra-subfield connectivity

We do not claim that our results represent the only patterns of communication between these anatomically complex regions. Moreover, while we investigated FC of broad portions of each subfield, we do not suggest that FC is segregated in such a coarse manner. Rather, the gradient nature of connectivity along hippocampal subfields is well documented in both the anatomical (Insausti and Munoz, 2001; Beaujoin et al., 2018) and functional (Vos de Wael et al., 2018; Maas et al., 2015; Libby et al., 2012) literatures (reviewed in Strange et al., 2014; Poppenk et al., 2013). In the current study, our method was not designed to investigate subtle intra-subfield gradients. Our rationale here was that, in line with the documented gradient nature of connectivity, different portions of each subfield would have a greater proportion of neurons functionally interacting with, for example, the cortical ROIs, and this would be reflected in a stronger correlation between their rsfMRI activity.

Our results complement previous studies which have observed functional gradients along the long axis of hippocampal subfields (Vos de Wael et al., 2018; Maas et al., 2015; Libby et al., 2012). It is important to note that within this handful of studies, a broad range of methods have been used to observe these functional gradients. While some have investigated hippocampal long axis connectivity in a cortex-wide manner (Vos de Wael et al., 2018), our study aimed to investigate in detail the FC of different portions of hippocampal subfields in relation to a small set of *apriori* ROIs. Also related to methodology, the placement of hippocampal subfield boundaries on MRI is an ongoing area of research yet to gain a consensus. Different hippocampal segmentation schemes have different criteria for subfield boundary placement (Yushkevich et al., 2015) and also incorporate different subfields in their segmentation protocols (Dalton et al., 2017; Berron et al., 2017). The results reported here pertain to the subfields as delineated according to the protocol by Dalton et al. (2017) and our results should be interpreted with these delineations in mind.

In conclusion, we have provided evidence that different hippocampal subfields have different patterns of FC with each other and with neighbouring cortical brain regions and, importantly, that this is also the case for different portions of each subfield. These differential patterns of FC may be facilitated, as we propose is the case for the association between the pre/parasubiculum and RSC, by anatomical connections which bypass the canonical hippocampal circuitry, permitting direct interactions to occur. This suggests that anterior and posterior portions of hippocampal subfields may be incorporated into different cortical networks which, in turn, might provide potential mechanisms by which functional differentiation along the long axis of the hippocampus can occur. Further investigations are required to assess the validity of this interpretation, and to investigate how these patterns of FC may be modulated by individual differences, normal ageing, in the context of brain pathologies and during the performance of different cognitive tasks.

## Acknowledgements

This work was supported by a Wellcome Principal Research Fellowship to E.A.M. (101759/Z/13/Z) and the Centre by a Centre award from Wellcome (203147/Z/16/Z).

The authors have no competing interests to declare.

## Abbreviations

A: Anterior (of the hippocampus)
AB: Anterior body (of the hippocampus)
CA 1-4: Cornu ammonis 1-4
DG: Dentate gyrus
DMN: Default mode network
ENT: Entorhinal cortex
FC: Functional connectivity
PB: Posterior body (of the hippocampus)
PHC: Posterior parahippocampal cortex
PRC: Perirhinal cortex
RSC: Retrosplenial cortex
rsfMRI: Resting state functional magnetic resonance imaging
T: Tail (of the hippocampus)

